# Incorporation of a Biocompatible Nanozyme in Cellular Antioxidant Enzyme Cascade Reverses Huntington’s Like Disorder in Preclinical Model

**DOI:** 10.1101/2020.09.23.310995

**Authors:** Aniruddha Adhikari, Susmita Mondal, Monojit Das, Pritam Biswas, Uttam Pal, Soumendra Darbar, Siddhartha Sankar Bhattacharya, Debasis Pal, Tanusri Saha-Dasgupta, Anjan Kumar Das, Asim Kumar Mallick, Samir Kumar Pal

## Abstract

The potentiality of nano-enzymes in therapeutic use has directed contemporary research to develop a substitute for natural enzymes, which are suffering from several disadvantages including low stability, high cost, and difficulty in storage. However, inherent toxicity, inefficiency in the physiological *milieu*, and incompatibility to function in cellular enzyme networks limit the therapeutic use of nanozymes in living systems. Here, we have shown that citrate functionalized manganese-based biocompatible nanoscale material (C-Mn_3_O_4_ NP) efficiently mimics glutathione peroxidase enzyme in the physiological *milieu* and easily incorporates into the cellular multienzyme cascade for H_2_O_2_ scavenging. A detailed computational study reveals the mechanism of the nanozyme action. We further established the *in vivo* therapeutic efficacy of C-Mn_3_O_4_ nanozyme in a preclinical animal model of Huntington’s disease, a prevalent progressive neurodegenerative disorder, which has no effective medication till date.

**SUMMARY:** Although, nano-enzymes have shown lots of promises in the management of several diseases, two major concerns limit their clinical translation. Apart from the inherent toxicity of the constituent materials (e.g., cerium, vanadium, gold, etc.), activities of contemporary nanozymes are often inhibited in physiological *milieu*. Furthermore, most of them are incapable of incorporation into the cellular metabolic networks for functioning in tandem and parallel with natural enzymes, a major criteria for potential therapeutics.

Here, we have shown that citrate-functionalized spherical Mn_3_O_4_ nanoparticles can efficiently mimic glutathione peroxidase (GPX) enzyme without the limitations of contemporary nanozymes, and effectively manage neurodegenerative Huntington’s disease in preclinical animal model. The choice of the material in the nanozyme lies on the fact that Mn is an essential micronutrient for mammals, and the stabilizing ligand citrate helps the nanoparticles to cross the blood-brain-barrier to reach brain. We have shown that the nanozyme can easily be incorporated in cellular antioxidant enzyme cascade. The specificity and efficacy of the nanozyme in the cascade was significantly higher compared to other reported nanozymes. We have justified our experimental findings with a detailed computational study. Understanding the mode of operation and management of Huntington’s disease in preclinical animal trial using a biocompatible (non-toxic) nanozyme as a part of the metabolic network may uncover a new paradigm in nanozyme based therapeutic strategy.

## MAIN TEXT

Over the past decade, nanozymes, nanomaterials with intrinsic enzyme-like properties have attracted significant interest for application in multiple fields owing to their advantages (i.e., high and tunable catalytic activity, low cost, easy large scale production, and high stability) over the drawbacks of natural enzymes (i.e., low stability, high cost, laborious preparation, and low recyclability) ^1-3^. Since the discovery of first iron-containing nanozyme in last decade ^4^, numerous nanomaterials have been elucidated to have oxidase ^5^, catalase ^6^, SOD ^7, 8^, peroxidase ^9^, monooxygenase ^10^, hydrolase ^11^, laccase ^12^ mimicking activities and therefore been used in diverse applications like the destruction of biofilm, removal of algal bloom, immunoassay, tissue staining, cancer treatment, and glucose biosensing. However, despite being the most promising candidate as catalytic biomedicine, the clinical translation of nanozymes for therapeutic usage is still lacking ^13-15^. Inherent toxicity of the materials used in the preparation of contemporary nanozymes, low aqueous solubility, inability to properly function in the physiological *milieu*, lack of selectivity (towards biological substrate), and incompatibility with other enzymes in catalyzing intracellular cascade reactions are considered to be the confounding factors ^15-18^

Highly connected networks of natural enzymes regulate the majority of the biological functions that occur in living systems. Dysregulation in any of these enzyme controlled networks often necessitates disease onset and progression. For example, redox imbalance due to downregulation of cellular antioxidant enzymes i.e., superoxide dismutase (SOD), catalase, or glutathione peroxidase (GPx) may lead to pathogenesis of cancer, diabetes, atherosclerosis, neurodegeneration, and aging ^19-22^. Recently, GSH dependent GPx with pan cellular distribution has emerged as the key antioxidant enzyme (presence of isoforms in both cytosol and mitochondria highlights its significance) for maintenance of cellular redox homeostasis ^23-25^. The deregulation in GPx activity and associated redox imbalance is associated with the pathogenesis of Huntington’s disease (HD), one of the most prevalent neurodegenerative disorders with early-onset and progressive fatality ^26-28^. Despite lack of effective therapeutics till date, one promising approach to treat the disease was found to be by replenishing the maladaptive enzyme (GPx) with an artificial one ^28^. However, to successfully introduce any artificial enzyme as a direct surrogate of traditional enzyme for therapeutic use, the cooperative functionality needs to be mimicked to allow cascade reactions to take place in a parallel and efficient manner ^29-31^. Recently, some of the nanozymes have been found to function in intracellular cascade reactions, however, concern over toxicity and metabolism have restricted their use in living organisms ^9, 32-34^. Thus, a biocompatible nanozyme that retains functionality in the physiological *milieu* and can easily be incorporated in the cellular enzymatic cascade is urgently needed for therapeutic usage.

Thus, considering the limitations and opportunities in clinical translation of nanozymes, particularly in neurodegenerative disorders, our objective for this study was to develop a non-toxic, aqueous soluble, biomimetic nanozyme capable of catalyzing intracellular cascade reactions and assess its therapeutic efficacy as redox medicine in an animal model of neurodegenerative disease where redox imbalance, associated oxidative distress and damage to the intracellular GPx system play a major role in the pathogenesis.

Here, we have shown that citrate functionalized Mn_3_O_4_ nanoparticles can efficiently mimic the enzymatic activity of glutathione peroxidase using GSH as co-factor. The nanozyme is highly specific towards H_2_O_2_ and can be incorporated into the glutathione reductase coupled reaction to scavenge H_2_O_2_, and oxidize NADPH simultaneously. Detailed experimental and computational studies reveal the mechanism of nanozyme action involving the generation of an intermediate peroxido species that accounts for the remarkable specific activity. Using, 3-nitro propionic acid (3-NPA) intoxicated C57BL/6j mice we confirmed that C-Mn_3_O_4_ NPs can cross the blood-brain barrier, retain its enzymatic activity in brain cells, and treat Huntington’s like neurodegenerative disorder in animals. Scavenging of intra and extramitochondrial ROS by GPx action, maintenance of cellular redox equilibrium, and subsequent reduction of oxidative damages lead to the therapeutic effect. Thus, the use of the C-Mn_3_O_4_ nanozyme as a nanomedicine against neurodegenerative disorders may uncover a new paradigm in nanozyme based therapeutic strategy.

Encouraged by the apparent non-toxicity (permissible limit ∼12 mg day^−1^), abundance of manganese (Mn) as the catalytic metal center or cofactor in several enzymes, and preferable e_g_ occupancy of 1.33 (*vide infra*) we selected nano-sized Mn_3_O_4_ as our compound of interest. A template or surfactant-free sol-gel based three-step approach was used to synthesize Mn_3_O_4_ nanoparticle at room temperature and pressure using MnCl_2_ as a precursor (detailed in Supplementary Materials and Methods). Citrate functionalization was performed to make the nanoparticle aqueous soluble, biocompatible, and competent to cross the blood-brain barrier (BBB) ^35, 36^. Transmission electron micrograph (TEM) shows the citrate functionalized Mn_3_O_4_ nanoparticles (C-Mn_3_O_4_ NPs) to be well-dispersed uniform spheres with an average diameter of ∼6.12±2.24 nm (Figure 1a & 1b). High resolution (HR) TEM image of a single nanoparticle confirms the crystalline nature with clear atomic lattice fringe spacing of 0.312±0.021 nm (Figure 1c) corresponding to the separation between (112) lattice planes. All x-ray diffraction (XRD) peaks corresponding to different planes of C-Mn_3_O_4_ NPs (Supplementary Figure S1a) accurately reflected the tetragonal hausmannite structure of Mn_3_O_4_ with lattice constants of a=5.76Å and c=9.47Å and space group of *I*41/amd as indexed in the literature (JCPDS No. 24-0734). Furrier transformed infrared (FTIR) spectra confirmed the binding of citrate to the surface of the nanomaterial (Supplementary Figure S1b). The hydrodynamic diameter of C-Mn_3_O_4_ NPs as measured using dynamic light scattering (DLS) was found to be ∼21.5±4.1 nm (polydispersity index, PDI ∼0.32) (Figure 1d) with zeta potential, ξ=-12.23±0.61 mV, and electrophoretic mobility −0.96±0.05 μcmV^−1^s.

**Figure 1.**
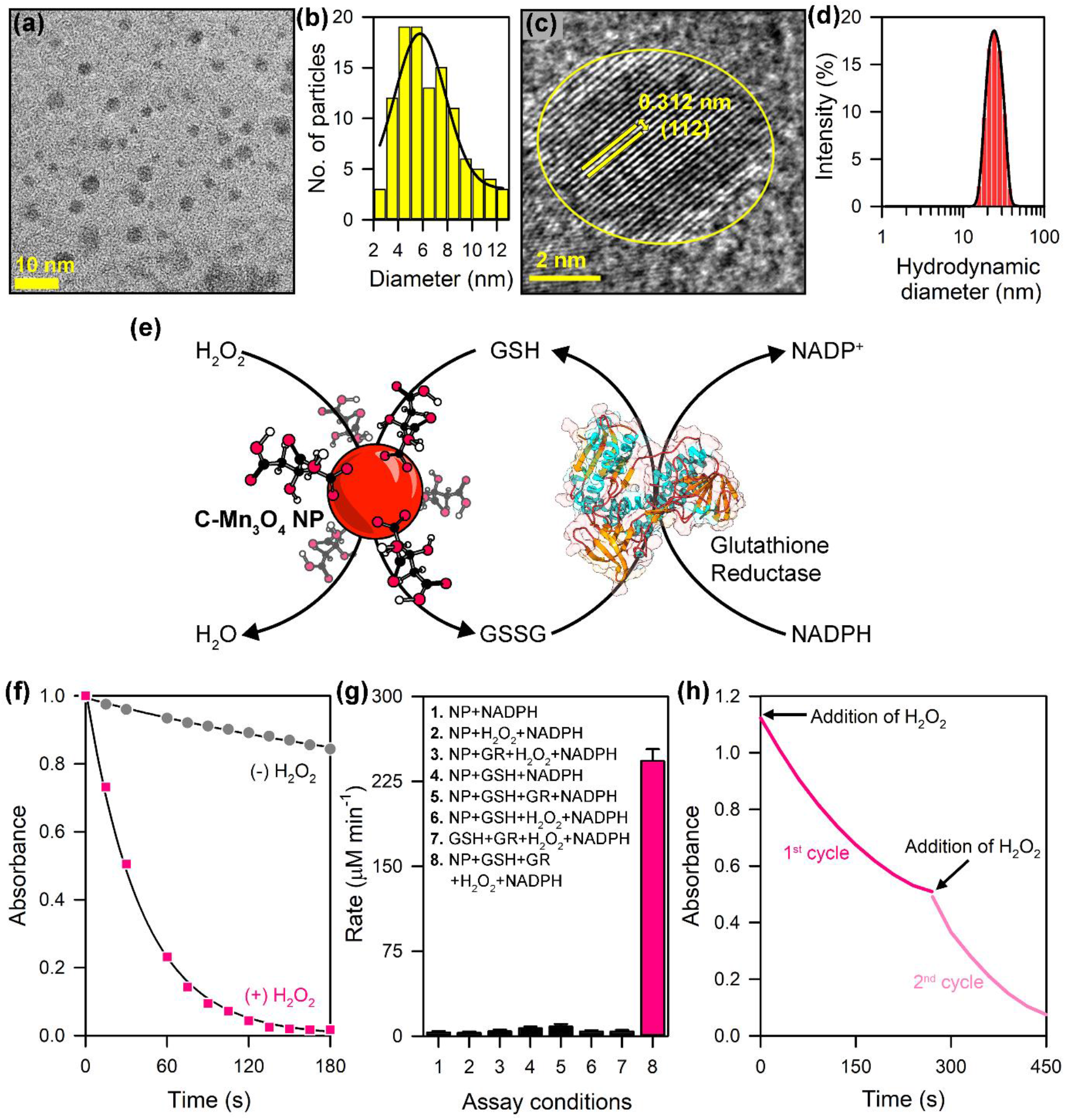
Characterization and GPx-mimetic activity of C-Mn_3_O_4_ NPs. (a) TEM image of C-Mn_3_O_4_ NPs. (b) Size distribution of the nanoparticles as measured from TEM. (c) HRTEM image of single nanoparticle. (d) Hydrodynamic diameter from DLS study. (e) Schematic diagram depicting the GPx-like activity of C-Mn_3_O_4_ NPs in presence of GSH, and recycling by GR in coupled reaction. (f) The change in absorbance of NADPH (340 nm) during the reaction. In absence of H_2_O_2_, no reactivity was observed. (g) Comparison of initial reaction rate at different assay conditions. (h) The reaction kinetics shows the mechanism to cyclic/ catalytic. The activity was studied for two cycles in UV-visible spectroscopy by addition of H_2_O_2_ (240 μM) and following the decrease of NADPH concentration at 340 nm.

The GPx mimetic activity of aqueous soluble C-Mn_3_O_4_ NP was evaluated at physiological pH (pH ∼7.4) using the glutathione reductase (GR)-coupled assay where the decrease in NADPH concentration was monitored spectrophotometrically at 340 nm. Figure 1e schematically illustrates the GPx like activity of C-Mn_3_O_4_ NPs in GR coupled cascade. Although nanozymes are known to have an incompatibility with other enzymes in a cascade ^2, 9^, we found the C-Mn_3_O_4_ NPs to be perfectly compatible with GR (Figure 1f). The reaction followed first-order reaction kinetics with a rate constant, *k* = 1.38±0.01 min^−1^ (Supplementary Figure S2a). The initial reaction rates determined at various assay conditions indicate that the GPx-like activity of C-Mn_3_O_4_ NPs is hindered by the absence of any one of the components in the reaction mixture (Figure 1g; Supplementary Figure S2b). The repeated H_2_O_2_ scavenging activity for several cycles indicates the reaction to be catalytic or recyclable (Figure 1h). A gradual increase in the initial reaction rate with an increasing concentration of C-Mn_3_O_4_ NPs was observed for the reduction of H_2_O_2_ (data not shown). The apparent steady-state kinetic parameters were determined by independently varying the concentrations of H_2_O_2_ (0-480 μM), and GSH (0-6.0 mM) in presence of GR (1.7 units), C-Mn_3_O_4_ NPs (1.3 μM), and NADPH (400 μM). Both reactions followed typical Michaelis-Menten kinetics (Figure 2a & 2c). The Michaelis-Menten constant (K_M_) and the maximum initial velocity (V_max_) were determined based on Lineweaver-Burk linearization (Figure 2b & 2d). K_M_ for H_2_O_2_ and GSH are ∼1.09±0.06 and ∼1.36±0.09 mM, respectively. V_max_ for H_2_O_2_ and GSH are ∼0.095±0.011 and ∼0.064±0.008 mM min^−1^, respectively. The K_M_ for H_2_O_2_ is higher compared to the natural GPx1 enzyme (∼0.01 mM) ^37, 38^ indicating a lower affinity towards substrate which is a very common phenomenon in artificial enzymes. Still, the observed K_M_ is lower compared to Ebselen (∼2.34 mM), the most studied GPx mimic; and equivalent to V_2_O_5_ nanowires (∼0.11 mM), one of the rare nanozymes that has the ability to be incorporated into enzymatic cascade but concern over toxicity limited their *in vivo* application. Interestingly, the affinity of C-Mn_3_O_4_ NPs for co-factor GSH is significantly higher compared to the native GPx1 (K_M_ ∼10 mM), indicating the explicit role of GSH in the catalytic activity of the nanozyme. The C-Mn_3_O_4_ nanozyme catalyzed the reduction of H_2_O_2_ with a turnover number (*k*_cat_) of ∼69.12±0.52 min^−1^ and an apparent second-order rate constant, *k*_cat_/K_M_ ∼10530.17±975.25 M s^−1^. For oxidation of GSH, the values were found to be *k*_cat_ ∼46.73±0.31 min^−1^ and *k*_cat_/K_M_ ∼5717.85±345.72 M s^−1^. Although, both turnover number and enzyme efficiency (for H_2_O_2_ reduction) were several orders of magnitude lower compared to the natural GPx1 enzyme isoform (*k*_cat_ ∼5780 min^−1^; *k*_cat_/K_M_ ∼9.63 × 10^6^ M s^−1^), C-Mn_3_O_4_ NPs outperformed several other artificial GPx mimics (Supplementary Table S1). For example, C-Mn_3_O_4_ NPs have shown ∼17 times higher turnover and ∼390 times higher enzyme efficiency compared to Ebselen (*k*_cat_ ∼3.85 min^−1^; *k*_cat_/K_M_ ∼27.33 M s^−1^); ∼18 times higher turnover and ∼17 times higher enzyme efficiency compared to V_2_O_5_ nanowires (*k*_cat_ ∼3.9 min^−1^; *k*_cat_/K_M_ ∼590 M s^−1^) (a detailed comparison with other GPx mimics is provided in Supplementary Table S1). As the kinetic data suggest, the remarkable enhancement in catalytic efficiency (*k*_cat_/K_M_) of C-Mn_3_O_4_ nanozyme was achieved not by reducing the K_M_ but through enhancement of the *k*_cat_, which is considered as one of the most challenging demands in the evolution of artificial enzymes ^39, 40^.

**Figure 2.**
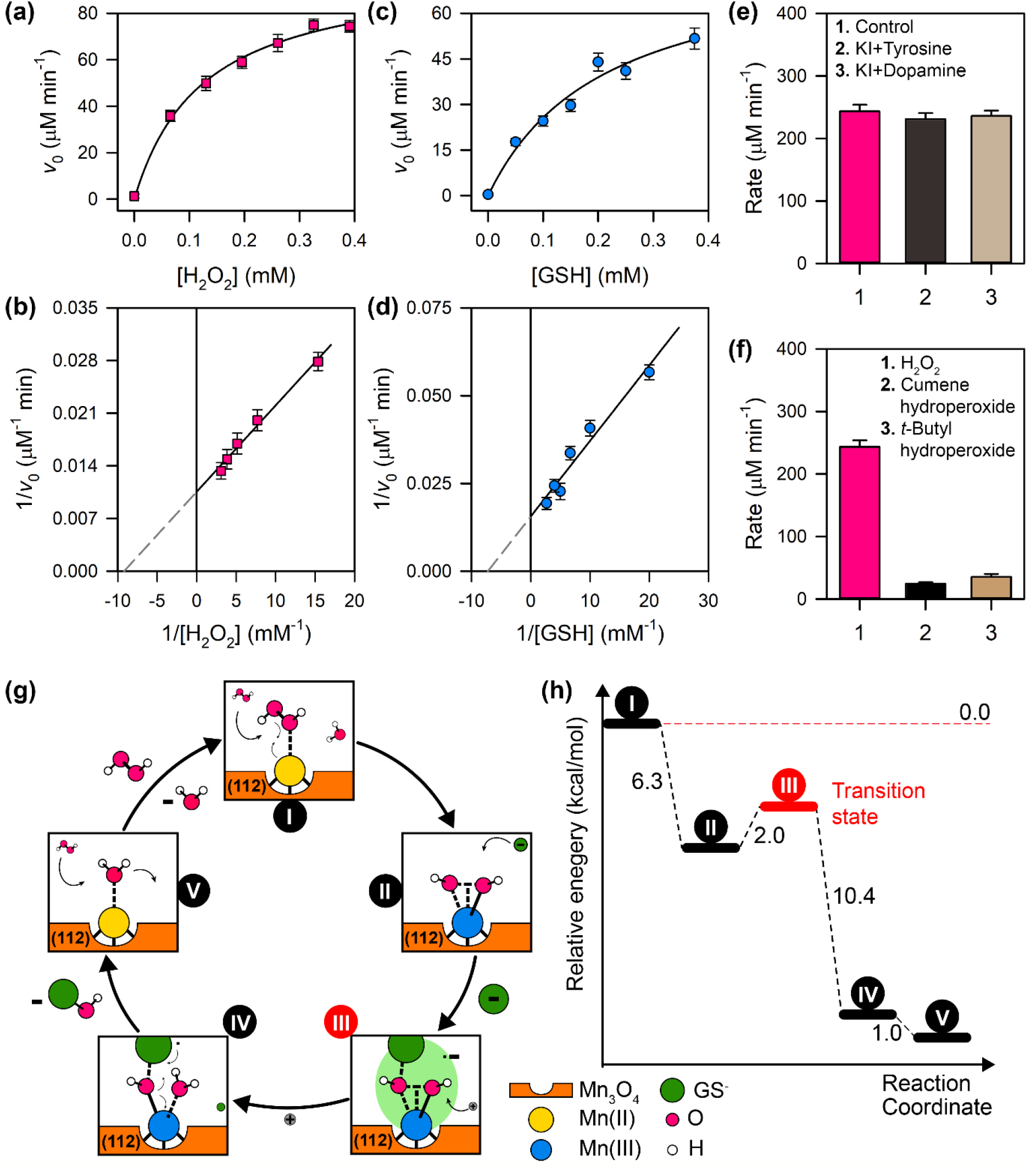
Kinetic parameters and mechanism of action. (a & b) Michaelis-Menten and Lineweaver-Burk plot for variable concentration of H_2_O_2_. (c &d) Michaelis-Menten and Lineweaver-Burk plot for variable concentration of GSH. (e) Effect of haloperoxidase substrate on GPx-like activity of C-Mn_3_O_4_ NP. (f) Selectivity of C-Mn_3_O_4_ NP towards H_2_O_2_. (g) Possible mechanism of GPx-like action of C-Mn_3_O_4_ NPs. (h) Energy profile for the reaction scheme.

Most of the peroxidase mimics exert their catalytic activity through the generation of •OH radical which in the presence of metal ions mediates the oxidation of organic substrates ^41, 42^. In order to investigate the role of •OH in the catalytic mechanism of C-Mn_3_O_4_ nanozyme, we introduced luminol, an •OH indictor into the reaction mixture. Non-appearance of any chemiluminescence signal indicated the absence of •OH during catalysis (Supplementary Figure S3). Therefore, the mechanism of C-Mn_3_O_4_ nanozyme was different from other peroxidase mimics. The exceptional selectivity of the nanozyme towards H_2_O_2_ is probably due to the formation of a polar peroxido species rather than •OH radicals ^9^. The peroxido species reacts further with the nucleophile (GSH cofactor) to form glutathione disulfide (GSSG) (detailed in computational studies). GPx mimics often tend to show haloperoxidase activity ^41^. So, we monitored the reaction of C-Mn_3_O_4_ NPs with H_2_O_2_ in the presence of haloperoxidase substrates i.e., dopamine/iodide or tyrosine/iodide. The reaction rate remained unaffected (Figure 2e). The unaltered reaction rate could be attributed to the facile attack of GS^−^ at the polarized oxygen atom of the peroxido species formed on the surface of C-Mn_3_O_4_ NPs upon reaction with H_2_O_2_ as a result of greater nucleophilic character of GS^−^ compared to halides. Further comparison between the reactivity of C-Mn_3_O_4_ NP (in terms of reaction rate) with various peroxide substrates e.g., H_2_O_2_, *t-*butyl hydroperoxide, and cumene hydroperoxide indicates that the catalytic action is specific to H_2_O_2_ (Figure 2f).

In order to validate the catalytic reaction mechanism of the nanozyme, as postulated above, a quantum chemical computational study using density functional theory (DFT) was performed (*vide* Supplementary Information for computational methods). The schematic of the reaction mechanism starting from the adsorption of H_2_O_2_ on the Mn(II) catalytic center to the formation of a water molecule and hydroxy-glutathione (GSOH) is illustrated in Figure 2g and Supplementary Figure S4a & S4b. The computed Gibbs free energy profile of the reaction path (Figure 2h) suggests that the adsorbed H_2_O_2_ spontaneously undergoes splitting (ΔG = −6.3 kcal/mol) on the Mn(II) center forming a peroxido species. In the next step, one proton is transferred from GSH to one of the OH groups attached to the catalytic center resulting in formation of water. This proton transfer process has a very low activation energy of 2 kcal/mol and a large Gibbs free energy of −10.4 kcal/mol suggesting a highly favourable reaction. After the proton donation, GS^−^ readily attacks the other OH group attached to the metal center and forms the GSOH intermediate, which then dissociates by regenerating the Mn(II) catalytic center. Water is then replaced by H_2_O_2_ and the next cycle begins. Thus, a •OH radical is never released. Water on the Mn(II) catalytic center is then replaced by H_2_O_2_ and the next cycle begins. GSOH undergoes a condensation reaction with a molecule of GSH to form the GSSG ^43^. Regarding the efficiency of such as catalysis, Wang et al. recently showed that e_g_ orbital (d_x2-y2_ and d_z2_) occupancy could be an excellent measure of peroxidase like activity of transition metal oxide nanozymes ^44^. Their study established a volcano relationship of activity with the average e_g_ occupancy on a salce of 0--2. i.e., the maximum activity was observed for a nanozyme with e_g_ occupancy of 1; the activity decreased as the e_g_ occupancy approaches 0 or 2. This explains why our Mn_3_O_4_ nanozyme (calculated e_g_ occupancy of ∼1.33) shows very high peroxidase like activity. In OH dissociation reaction during water oxidation as well, Saha-Dasgupta et al. previously showed that population of e_g_ state in the high spin Mn catalytic center of Mn_4_O_4_ cubane is associated with their higher catalytic efficiency over other transition metal catalysts such as Co_4_O_4_ ^45^. The mechanism is illustrated in Figure 2g and Supplementary Figure S4. The energy diagram for different steps is indicated in Figure 2h.

Preclinical animal studies are essential for the translation of potential therapies from bench to bedside ^46^. Considering the *in vitro* adaptability of C-Mn_3_O_4_ nanozyme in redox regulatory mechanisms we tested their efficacy in an animal model as a prelude to clinical translation. Huntington’s disease (HD), one of the most prevalent neurodegenerative disorders, is an autosomal-dominant disorder caused by an expansion of CAG repeats in the gene huntingtin, *htt*, and characterized by lesions in the striatum of the brain that cause progressive behavioral and cognitive impairments and involuntary choreiform movements ^26, 27^. Unfortunately, to date, there is no satisfactory medicine to prevent or slow the pathogenesis of HD ^27, 47^. Strong evidence suggests a causal relationship between oxidative stress and HD ^27^. Elevated markers of oxidative damage such as protein oxidation, lipid peroxidation, and DNA damage have been linked to the pathogenesis. Other than oxidative stress, transcriptional impairment, excitotoxicity, inflammation, apoptosis, and mitochondrial dysfunction leads to disease onset and striatal degeneration ^48^. Interestingly in one recent study, Mason et. al., has shown that overexpression of GPx (neither SOD nor catalase) in cellular, yeast, or drosophila models of HD could mitigate the mHtt (mutant huntingtin) toxicity and associated oxidative damage ^28^. Therefore, we hypothesized that pharmacological interventions with GPx mimic like C-Mn_3_O_4_ nanozyme could be a viable treatment option to prevent HD pathogenesis considering the mimic would be able to cross the BBB, enter the brain cells and efficiently supplement the intracellular GPx activity. We selected a well-studied 3-NPA induced C57BL/6j mice model of HD to test the *in vivo* therapeutic efficacy of C-Mn_3_O_4_ nanozyme ^49-52^. 3-NPA is known to inhibit mitochondrial respiratory complex-II (succinate dehydrogenase, SDH) in neuronal cells instigating mitochondrial impairment, ATP depletion, increase in reactive oxygen species (ROS), excitotoxicity and thereby, simulates a neurobehavioral condition exactly similar to mHtt toxicity and HD ^53^.

Several studies have indicated that 3-NPA damages striatal medium spiny neurons, which lead to a progressive deficit in fine and gross motor function ^51^. Motor function was evaluated through four tests: beam traversal, pole descent, nasal adhesive removal, and hindlimb clasping reflexes. 3-NPA treatment caused a progressive decline in the hind limb clasping reflex score, a hallmark of HD pathogenesis, and definitive measure of striatal dysfunction throughout the experimental regime (Figure 3a). Treatment with C-Mn_3_O_4_ nanozymes protected the animals from derogatory motor impairment. From the 10^th^ day of the experimental period the hind limb clasping reflex started to improve significantly compared to the 3-NPA intoxicated group (Figure 3a). The other two groups (control and C-Mn_3_O_4_ nanozyme treated) retained baseline performance. Treatment solely with the constituent ligand citrate did not affect clasping behavior (Supplementary Figure S5a). 3-NPA-administered mice required significantly extra time to cross a challenging beam (Figure 3b), and to descend a pole (Figure 3c), the two methods of accessing gross motor function, compared to untreated control or C-Mn_3_O_4_ nanozyme-treated littermates. Treatment with C-Mn_3_O_4_ nanozyme resulted in significant improvement in both time to cross a challenging beam, and to descend a pole (Figure 3b & 3c). In contrast, treatment with citrate showed no signs of improvement in the tests (Supplementary Figure S5). Removal of an adhesive from the nasal bridge, which provides information about fine motor control, was impaired in 3-NPA intoxicated mice compared to the other three groups (Figure 3d). While, citrate treated animals displayed similar results to 3-NPA treated ones (Supplementary Figure S5d). The observed recovery in both gross motor function, as well as fine motor control due to treatement with C-Mn_3_O_4_ nanozyme was found to be dose-dependent (Supplementary Figure S5). Considering the therapeutic inefficacy of citrate, we excluded the citrate treated group from further experiments. The striatum and related basal ganglia circuits are known to contribute towards the acquisition of repetitive and stereotyped behaviors ^54, 55^. To acess fine motor movements, we performed rotarod test. In the rotarod study, motor learning was evaluated considering the improvement in performance (i.e., latency to fall) over three trials. In 3-NPA-treated mice, latencies to fall were lesser in delayed tests at day-1 and day-15 (Figure 3e), which is a marker of fine-motor function deficit. For the group that received both 3-NPA and C-Mn_3_O_4_ NPs, latency to fall was significantly increased compared to the 3-NPA treated group. So, the C-Mn_3_O_4_ nanozyme was successful in improving the fine motor movements casued by 3-NPA administration. To evaluate the sensory motor functions, we used tail-flick assay (Figure 3f), where the 3-NPA intoxicated mice in the first trial exhibited lengthier tail-flick latency, thus poorer pain sensitivity. However, this variance was not observed in successive trials (data not shown). The tail-flick latencies of 3-NPA+C-Mn_3_O_4_ NP treated, and C-Mn_3_O_4_ NP treated groups were similar to that of untreated control (Figure 3f). To be sure, we compiled all motor phenotypes into a principal component analysis (PCoA). The result displays the prominent segregation of the 3-NPA intoxicated group with the others (Figure 3g). The animals co-treated with 3-NPA+C-Mn_3_O_4_ NPs, or C-Mn_3_O_4_ NPs alone clusped together with the control animals (Figure 3g). Collectively, these results indicate that C-Mn_3_O_4_ nanozyme significantly protected 3-NPA intoxicated mice from the hallmark motor dysfunctions that resemble HD-like syndrome.

**Figure 3.**
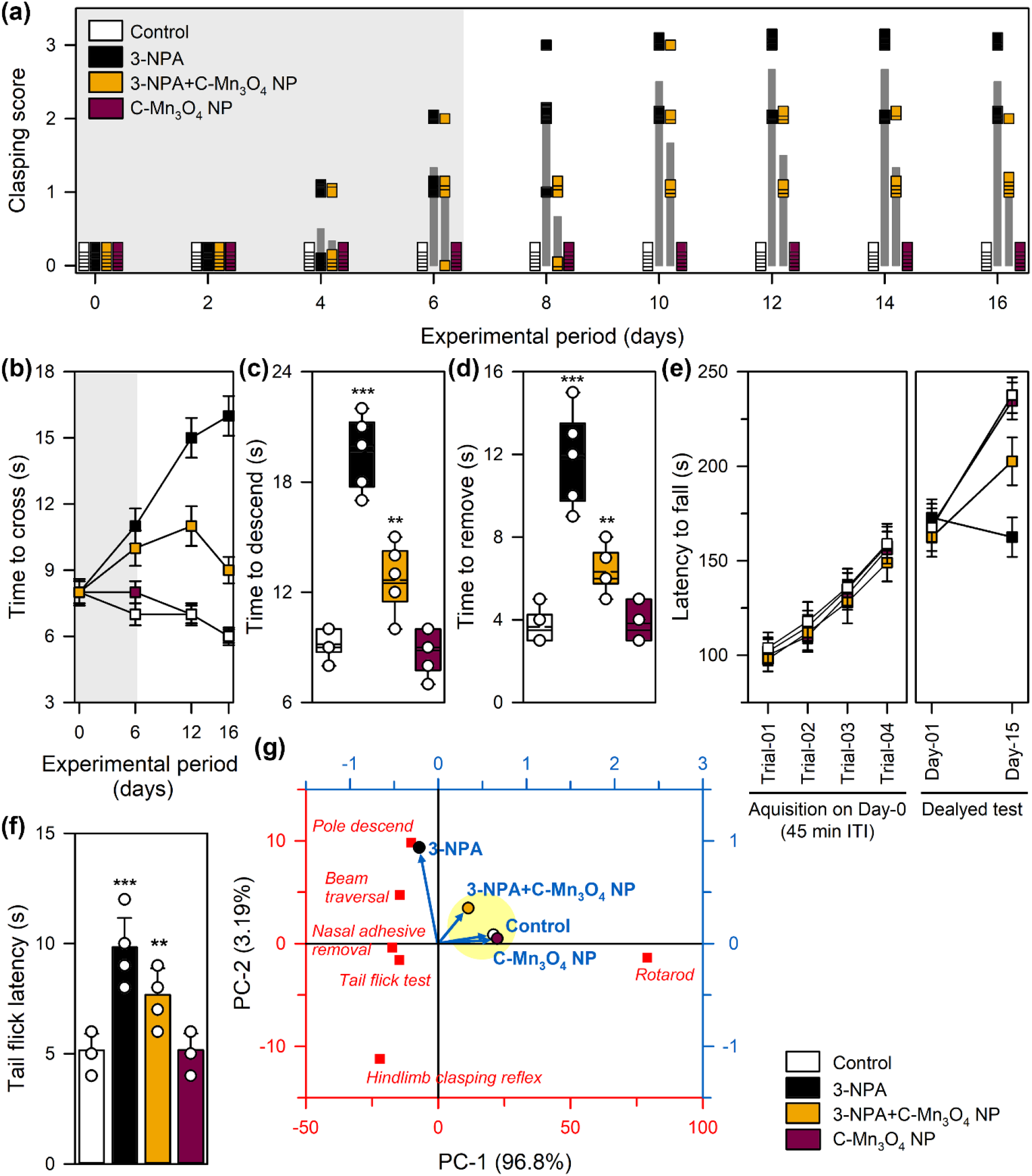
Effect of C-Mn_3_O_4_ NPs on 3-NPA induced motor impairment, hallmark of Huntington’s disorder. (a) Hind limb clasping reflex shows progressive improvement due to C-Mn_3_O_4_ NP treatment during the experimental regime. Darker shaded region represents co-treatment period. (b) Time to cross a beam in beam traversal test. (c) Time to descend a pole. (d) Nasal adhesive removal time. (e) Rotarod test. (f) Tail flick latency. (g) PCA analysis considering all motor phenotypes. Data are expressed as Mean ± SD. N= 6. *, **, *** Values differ significantly from control group (without treatment) (***p< 0.001; **p< 0.01; *p< 0.05).

Previous studies indicate that 3-NPA causes striatal damage and produces anxiety-like behavior, similar to human HD ^52, 56^. Therefore, we evaluated the effect of C-Mn_3_O_4_ NPs on the anxiolytic behavior of 3-NPA induction. The thigmotactic behavior as an index of anxiety of the animals were evaluated using open field test (OFT). 3-NPA intoxication significantly enhanced the thigmotactic behavior (an indicator of increased anxiety), as showed by the lesser affinity of the animals to spent time in the central zone of the apparatus (Figure 4a-4b). Animals solely treated with C-Mn_3_O_4_ nanozyme spent the highest time at the central zone, while 3-NPA+C-Mn_3_O_4_ nanozyme-treated animals spent sufficiently enough time compared to control animals. The total distance moved by the animals and velocity were similar for all the groups, with the exception of the 3-NPA intoxicated mice (Figure 4c & 4d). The deteriorated locomotor function was probably one of the major reasons behind this observation. Nevertheless, it can reasonably indicate that the C-Mn_3_O_4_ nanozymes possess anxiolytic property that commendably overturned the 3-NPA induced anxiety-like behavior. To further check this hypothesis, elevated plus maze (EPM) tests for anxiety-like behavior was employed. Consistent with previous studies, the 3-NPA-intoxicated mice spent significantly lesser amount of time in the open arms of the EPM compared to the other three groups (Figure 4e and 4f). Similar observations were found in terms of the distance they moved in the open arms (Figure 4g). Similar to OFT, in case of EPM too, the total distance moved was lesser for the 3-NPA-treated mice (Figure 4h). In agreement with this behavior, the 3-NPA-treated mice spent more time in the closed arms of the apparatus. Both OFT and EPM studies were performed in a regular time interval, throughout the experimental period. The results showed similar trends like other motor functions discussed in previous section (data not shown). Combination of the observed behavioral features indicated that the anxiety was induced due to severe 3-NPA neurotoxicity, which was ameliorated upon treatment with C-Mn_3_O_4_ nanozymes. Light preference test was further used to validate our observations about the anxiolytic effects of C-Mn_3_O_4_ nanozymes. In light preference test, reduced movement in the light area is considered as an indicator of anxiety. 3-NPA-treated mice exhibited both lesser activity and transitions in the light zone of the apparatus, while treatment with C-Mn_3_O_4_ nanozyme efficiently recovered their normal activity (Figure 4i).

**Figure 4.**
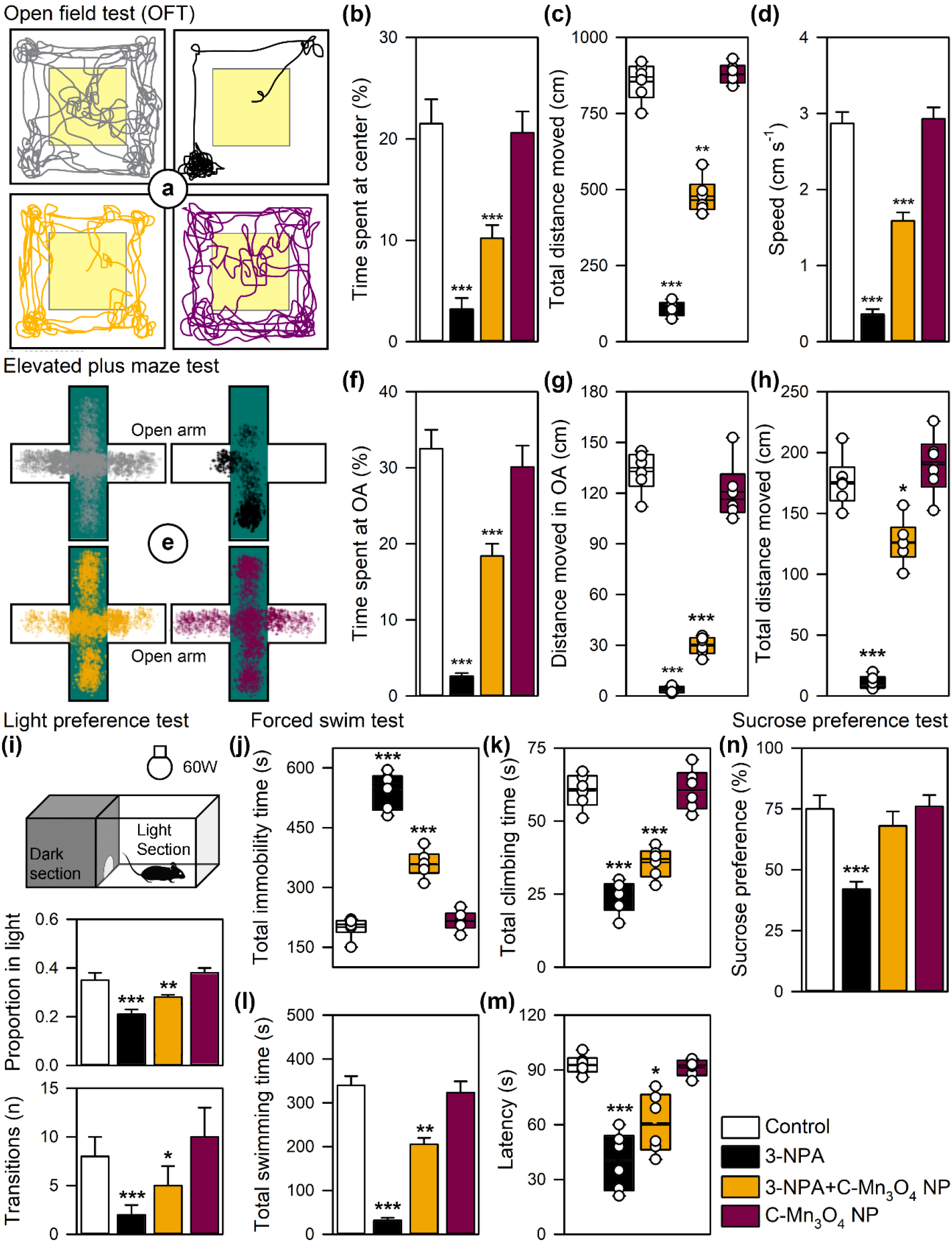
Effect of C-Mn_3_O_4_ NPs on anxiety and depression-like behavior. Open field test. (a) Trace of open field activity. (b) Time spent at the center. (c) Total distance moved. (d) Average speed. Elevated plus maze test. (e) Trace of movement in EPM. (f) Time spent in open arm. (g) Distance moved in open arm. (h) Total distance moved. (i) Light preference test. Time spent in light zone and transitions into light zone. Forced swim test. (j) Total immobility time, (k) total climbing time, (l) total swimming time and (m) latency to first immobility event. (n) Sucrose preference test. Data are expressed as Mean ± SD. N= 6. *, **, *** Values differ significantly from control group (without treatment) (***p< 0.001; **p< 0.01; *p< 0.05).

Another prominent feature of 3-NPA-induced neurotoxicity is the introduction of depression-like behavior in rodents, a symptom similar to the HD affected human counterpart. Therefore, we employed the forced swim test (FST) to assess depression-like behavior. In FST, high immobility time reflects increased depression. Consistent with previous studies, 3-NPA treatment resulted in increased immobility time (Figure 4j). Treatment with C-Mn_3_O_4_ nanozyme reduced the indications of depression as imitated in the lower immobility time (Figure 4j). The untreated control group and C-Mn_3_O_4_ nanozyme treated group showed comparable results. Measurements of the climbing time, swimming time, and latency to the first immobility event further highlighted the antidepressant-like action of the nanozyme. The climbing activity was significantly lower in 3-NPA-treated animals (Figure 4k). There was no observable improvement even after treatment with the nanozyme. Total swimming time (Figure 4l), or latency to first immobility (Figure 4m) were comparable throughout all four groups. Observed depression-like behavior of the 3-NPA-treated mice was accompanied by anhedonia (i.e., inability to experience pleasure from activities usually found enjoyable, in this case tasting the sweetness of sucrose), as indicated by the sucrose preference test (SPT) (Figure 4n). The preference for sucrose was almost identical for all the other three groups, demonstrating the healing effect of the C-Mn_3_O_4_ nanozyme (Figure 4n).

According to previous studies, 3-NPA exposure may increase oxidative stress in the hippocampus leading to memory deficit and affective disturbances reminiscent of the HD ^57, 58^. To characterize whether C-Mn_3_O_4_ nanozyme can reverse the 3-NPA persuaded hippocampal damage, we employed novel object recognition, and Morris water maze (MWM) tests. Novel obeject recognition was used to illuminate the behavioral complications (disturbances in recall memory) due to 3-NPA-intoxication, and the effect of the nanozyme over it. Next to three days of 15 min habituation trials in the testing apparatus, animals were permitted to explore two identical objects for 5 min and were then returned to their home cages. After an interval of 60 mins, one familiar object was replaced with a novel object, and the animals were permitted another 2 mins of exploration time. Their time of interaction with each of the two objects were measured during the experimental period (Supplementary Figure S6a). As anticipated, the untreated control animals with entirely intact recall memory, spent more time with the novel object than the familiar one (Supplementary Figure S6b and S6c). In contrast, 3-NPA-intoxicated animals were unable to discriminate between the novel and familiar objects, and spent almost similar time with both of them (Supplementary Figure S6b and S6c). Conversely, treatment with C-Mn_3_O_4_ nanozymes recovered the 3-NPA-treated animals from the profound cognitive deficit resulted from disturbances in hippocampus dependent learning and memory (Supplementary Figure S6b and S6c). To further verify the results of novel object recognition, MWM test was used. In case of 3-NPA-intoxicated animals, severe declines in spatial learning was found as the animals were failed to find the platform within provided timeframe (Supplementary Figure S7a). The amount of time the 3-NPA treated animals spent in the target quadrant was nominal and insignificant (Supplementary Figure S7b and S7c). In contrst, the animals co-treated with 3-NPA and C-Mn_3_O_4_ nanozyme were successful in finding the hidden platform, although the time taken to reach the platform was longer compared to the control animals (Supplementary Figure S7b and S7c). The other group, i.e.the C-Mn_3_O_4_ nanozyme-treated group, performed similar to the untreated control group (Supplementary Figure S7b and S7c). Results of the aforementioned two tests indicate towards severe hippocampal damage caused due to the 3-NPA treatment, and its prevention by C-Mn_3_O_4_ nanozyme treatment.

The results of behavioural studies were further supported by our histopathological findings (Figure 5). The hematoxylin and eosine stained brain sections of control mice showed normal brail tissue architecture. In 3-NPA treated mice, several signs of damage, particularly increase in apoptotic cells were evident in cerebellum and basal ganglia region of the brain. In basal ganglia focal degeneration of cells were also observed. In cerebellum region, the number of Purkinje cells were found to be reduced. Fibrillary gliosis was also evident in some regions. In co-treated (3-NPA+ C-Mn_3_O_4_ NP) and C-Mn_3_O_4_ NP-treated mice no significant damage in brain cell architechture was observed. This clearly indicated that treatment with the nanozyme decreased the Huntington like damage in the brain. It further implies that, the NPs are extremey safe to administer for treatment of neuronal damages.

**Figure 5.**
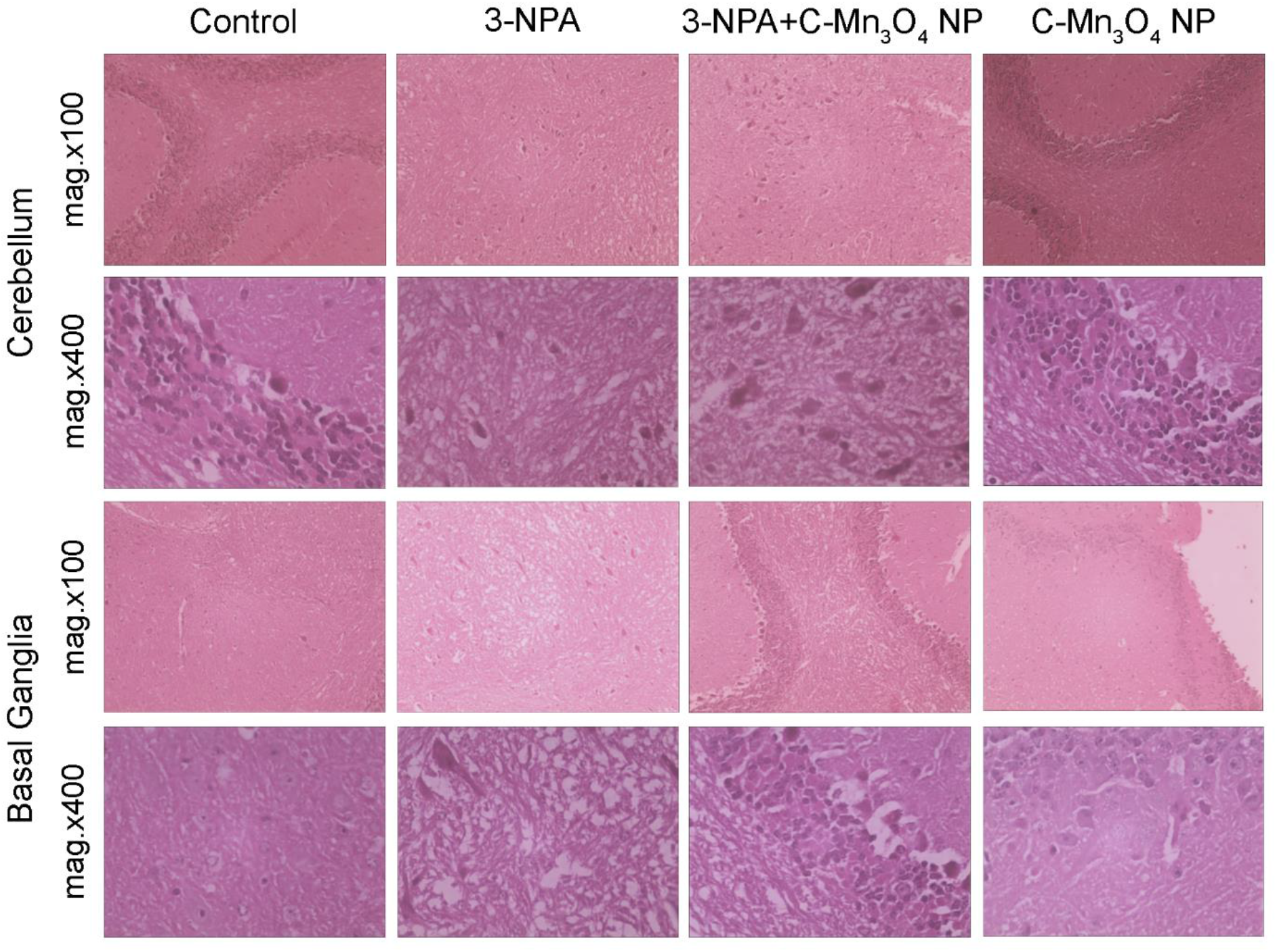
Effect of C-Mn_3_O_4_ NPs on 3-NPA induced histopathological damages. Microscopic images of haematoxylin and eosin stained sections of cerebellum and basal ganglia are shown at 100X and 400X magnifications.

To changes in the behavioral phenotype and morphometric histological findings indicate toards the retention of GPx mimic activity *in vivo*. To further confirm its effects, we measured the lipid peroxidation in the brain tissue. Upon 3-NPA administration the lipid peroxidation increased significantly, while the NPs were able to protect the neuronal cells from its. The SOD, catalase and GPx activities were almost identical from the two groups, while less lipid peroxidation indicated towards the GPx mimic activity of C-Mn_3_O_4_ NPs. As the oxidative damage decreased, it was expected that the mitochondrial damage will decline upon nanozyme treatment. In rodents, high doses of 3-NPA cause degeneration of striatal neurons and motor dysfunction similar to Huntington’s disease ^59^. The primary mechanism of 3-NPA-induced neurotoxicity involves the suicide inhibition of the mitochondrial electron transport chain (ETC) linked enzyme succinate dehydrogenase (SDH or complex-II) ^49^. Inhibition of SDH interferes with ETC and oxidative phosphorylation leading to cellular energy deficit (a decrease in ATP production) ^50^, oxidative stress, depletion of reduced glutathione (GSH), and alteration in the activities of cellular antioxidant enzymes ^53^. In agreement with previous observations, 3-NPA alone increased the lipid peroxidation (Figure 6a), a marker of oxidative damage, and significantly reduced the activities of cellular antioxidant enzymes, SOD, Catalase, and GPx (Figure 6b-6d). Treatment with C-Mn_3_O_4_ NPs significantly increased the activity of GPx (Figure 6d) and to some extent rescued the activities of SOD and catalase (Figure 6b & 6c). The observed change in GPx activity can be attributed to the GPx-mimic activity of the nanozyme to support the H_2_O_2_ scavenging by natural GPx. Whereas, the regain of SOD and catalase activities may be due to the indirect beneficial effect of an overall decrease in oxidative distress reflected in the reduction in lipid peroxidation (Figure 6a). Consistent with previous observations, in the current *in vivo* model of HD, we found that brain mitochondrial function was impaired in 3-NPA-treated animals (Figure 6e-6j). Increased mitochondrial permeabilization (mitochondrial swelling or mPTP formation) (Figure 6e), deregulated mitochondrial membrane potential (Figure 6f), decreased ATP level (Figure 6g), decreased mitochondrial dehydrogenase activity (Complex-II; Figure 6h), decreased complex IV activity (Figure 6i), increased mitochondrial ROS (Figure 6j) were evident in the brain tissue of HD animals. These deteriorating changes in mitochondrial parameters resulted in neuronal degeneration observed in the histological findings and motor behaviours. C-Mn_3_O_4_ NPs were able to efficiently protect mitochondria from the aforementioned damage (Figure 6e-6j). Our results strongly suggest that the GPx mimic activity of C-Mn_3_O_4_ nanozyme helped in scavenging the free radicals and reducing the associated oxidative damage, thereby prevented the mitochondrial dysfunctions and concomitant redox imbalance, the major underlying cause of neurodegenerative diseases like HD.

**Figure 6.**
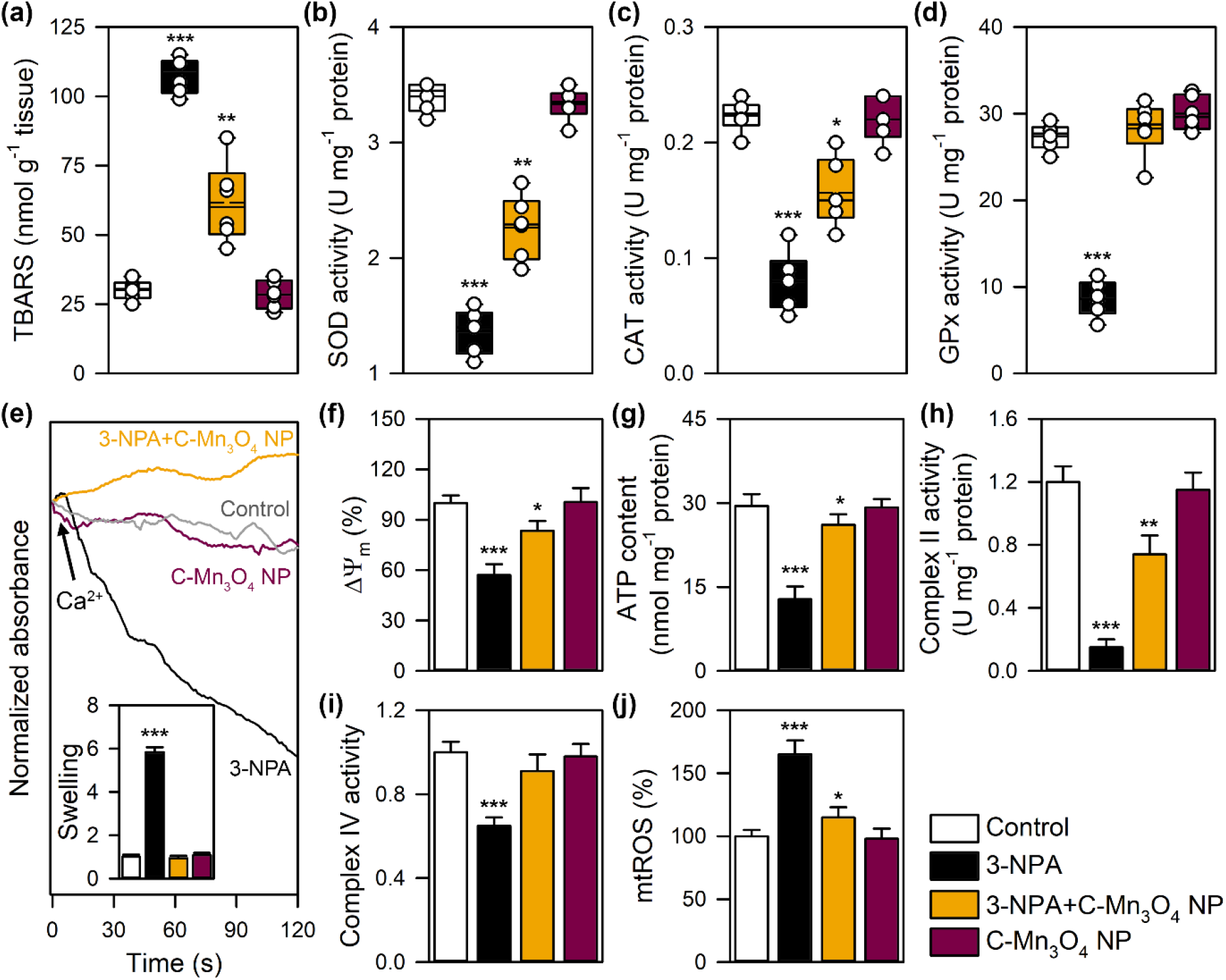
Effect of C-Mn_3_O_4_ NPs on antioxidant enzymes and mitochondrial parameters of brain. (a) Lipid peroxidation. (b) SOD activity. (c) Catalase activity. (d) GPx activity. (e) Effect on mitochondria permeability transition, measured as a decrease in absorbance at 540 nm. Inset shows extent of swelling in comparison to the control. (f) Change in mitochondrial membrane potential (ΔΨ_m_). (g) ATP content. (h) Complex-II activity of the electron transport chain. (i) Complex-IV activity of the electron transport chain. (j) Mitochondrial ROS (mtROS) as measured using DCFH assay. Data are expressed as Mean ± SD. N= 6. *, **, *** Values differ significantly from control group (without treatment) (***p< 0.001; **p< 0.01; *p< 0.05).

## CONCLUSION

In conclusion, our study suggests that in neutral pH and temperature (or, in physiological milieu) C-Mn_3_O_4_ NPs possess distinctive GSH dependent GPx mimic activity with excellent catalytic efficiency and substrate selectivity, the two most important parameters of evaluating artificial enzymes for therapeutic use essentially in a non-toxic manner. The unique ability of the nanozyme to be incorporated in a cellular redox modulatory enzyme network without causing adverse side effects makes C-Mn_3_O_4_ NPs suitable for biomedical application. A detailed computational study reveals mechanistic pathway behind the enzymatic action and is consistent with the *in vitro* studies. Results of animal studies further demonstrate its ability to pass the blood-brain barrier and treat progressive neurodegenerative disorders like Huntington’s disese. Enzymatic scavenging of neuronal reactive oxygen species and subsequent protection of mitochondria from oxidative damage resulted in the observed therapeutic effect. Successful clinical translation may open a new avenue in the nanozyme bases therapeutics of several other diseases (e.g., cancer, diabetes, Parkinson’s, Alzheimer’s, cardiovascular and chronic kidney diseases) where redox imbalance plays a key role in the pathogenesis.

## ACKNOWLEDGMENTS

MD thanks University Grants Commission (UGC), Govt. of India for Junior Research Fellowship. SKP thanks the Indian National Academy of Engineering (INAE) for the Abdul Kalam Technology Innovation National Fellowship, INAE/121/AKF. The authors thank the DBT (WB)-BOOST scheme for the financial grant, 339/WBBDC/1P-2/2013.

## REFERENCES

1. M. Liang and X. Yan, Accounts of Chemical Research, 2019, 52, 2190–2200.

2. H. Wei and E. Wang, Chemical Society Reviews, 2013, 42, 6060–6093.

3. B. Jiang, D. Duan, L. Gao, M. Zhou, K. Fan, Y. Tang, J. Xi, Y. Bi, Z. Tong, G. F. Gao, N. Xie, A. Tang, G. Nie, M. Liang and X. Yan, Nature Protocols, 2018, 13, 1506–1520.

4. L. Gao, J. Zhuang, L. Nie, J. Zhang, Y. Zhang, N. Gu, T. Wang, J. Feng, D. Yang, S. Perrett and X. Yan, Nature Nanotechnology, 2007, 2, 577–583.

5. L. Jiang, S. Fernandez-Garcia, M. Tinoco, Z. Yan, Q. Xue, G. Blanco, J. J. Calvino, A. B. Hungria and X. Chen, ACS Applied Materials & Interfaces, 2017, 9, 18595–18608.

6. W. Zhang, S. Hu, J.-J. Yin, W. He, W. Lu, M. Ma, N. Gu and Y. Zhang, Journal of the American Chemical Society, 2016, 138, 5860–5865.

7. R. Ragg, A. M. Schilmann, K. Korschelt, C. Wieseotte, M. Kluenker, M. Viel, L. Völker, S. Preiß, J. Herzberger, H. Frey, K. Heinze, P. Blümler, M. N. Tahir, F. Natalio and W. Tremel, Journal of Materials Chemistry B, 2016, 4, 7423–7428.

8. G. Wu, V. Berka, P. J. Derry, K. Mendoza, E. Kakadiaris, T. Roy, T. A. Kent, J. M. Tour and A.-L. Tsai, ACS Nano, 2019, 13, 11203–11213.

9. A. A. Vernekar, D. Sinha, S. Srivastava, P. U. Paramasivam, P. D’Silva and G. Mugesh, Nature Communications, 2014, 5, 5301.

10. B. E. R. Snyder, P. Vanelderen, M. L. Bols, S. D. Hallaert, L. H. Böttger, L. Ungur, K. Pierloot, R. A. Schoonheydt, B. F. Sels and E. I. Solomon, Nature, 2016, 536, 317–321.

11. M. Sun, L. Xu, A. Qu, P. Zhao, T. Hao, W. Ma, C. Hao, X. Wen, F. M. Colombari, A. F. de Moura, N. A. Kotov, C. Xu and H. Kuang, Nature Chemistry, 2018, 10, 821–830.

12. H. Liang, F. Lin, Z. Zhang, B. Liu, S. Jiang, Q. Yuan and J. Liu, ACS Applied Materials & Interfaces, 2017, 9, 1352–1360.

13. P. Wang, S. Liu, M. Hu, H. Zhang, D. Duan, J. He, J. Hong, R. Lv, H. S. Choi, X. Yan and M. Liang, Advanced Functional Materials, n/a, 2000647.

14. S. Hong, Q.-L. Zhang, D.-W. Zheng, C. Zhang, Y. Zhang, J.-J. Ye, H. Cheng and X.-Z. Zhang, iScience, 2020, 23, 100778.

15. D. P. Cormode, L. Gao and H. Koo, Trends in Biotechnology, 2018, 36, 15–29.

16. Y. Lv, M. Ma, Y. Huang and Y. Xia, Chemistry – A European Journal, 2019, 25, 954–960.

17. K. Fan, H. Wang, J. Xi, Q. Liu, X. Meng, D. Duan, L. Gao and X. Yan, Chemical Communications, 2017, 53, 424–427.

18. Y. Yoshihisa, Q.-L. Zhao, M. A. Hassan, Z.-L. Wei, M. Furuichi, Y. Miyamoto, T. Kondo and T. Shimizu, Free Radical Research, 2011, 45, 326–335.

19. P. Storz, Science’s STKE, 2006, 2006, re3–re3.

20. T. Finkel, Nature Reviews Molecular Cell Biology, 2005, 6, 971–976.

21. T. Finkel and N. J. Holbrook, Nature, 2000, 408, 239–247.

22. M. Patel, Trends in Pharmacological Sciences, 2016, 37, 768–778.

23. J. R. Arthur, Cellular and Molecular Life Sciences, 2001, 57, 1825–1835.

24. R. Brigelius-Flohé, Free Radical Biology and Medicine, 1999, 27, 951–965.

25. R. Brigelius-Flohé and M. Maiorino, Biochimica et Biophysica Acta (BBA) - General Subjects, 2013, 1830, 3289–3303.

26. J. K. McGill and M. F. Beal, Cell, 2006, 127, 465–468.

27. B. D. Paul and S. H. Snyder, Frontiers in Molecular Neuroscience, 2019, 12.

28. R. P. Mason, M. Casu, N. Butler, C. Breda, S. Campesan, J. Clapp, E. W. Green, D. Dhulkhed, C. P. Kyriacou and F. Giorgini, Nature Genetics, 2013, 45, 1249–1254.

29. J. E. Dueber, G. C. Wu, G. R. Malmirchegini, T. S. Moon, C. J. Petzold, A. V. Ullal, K. L. J. Prather and J. D. Keasling, Nature Biotechnology, 2009, 27, 753–759.

30. B. Wörsdörfer, K. J. Woycechowsky and D. Hilvert, Science, 2011, 331, 589–592.

31. J. D. Keasling, ACS Chemical Biology, 2008, 3, 64–76.

32. J. M. Wörle-Knirsch, K. Kern, C. Schleh, C. Adelhelm, C. Feldmann and H. F. Krug, Environmental Science & Technology, 2007, 41, 331–336.

33. E.-J. Park, G.-H. Lee, C. Yoon and D.-W. Kim, Environmental Research, 2016, 150, 154–165.

34. W. Lin, Y.-w. Huang, X.-D. Zhou and Y. Ma, International Journal of Toxicology, 2006, 25, 451–457.

35. R. A. Yokel, NeuroMolecular Medicine, 2009, 11, 297–310.

36. R. A. Yokel, M. Wilson, W. R. Harris and A. P. Halestrap, Brain Research, 2002, 930, 101–110.

37. L. Flohé and I. Brand, Biochimica et Biophysica Acta (BBA) -Enzymology, 1969, 191, 541–549.

38. D. E. Paglia and W. N. Valentine, The Journal of laboratory and clinical medicine, 1967, 70, 158–169.

39. R. Obexer, A. Godina, X. Garrabou, P. R. E. Mittl, D. Baker, A. D. Griffiths and D. Hilvert, Nature Chemistry, 2017, 9, 50–56.

40. W. J. Albery and J. R. Knowles, Biochemistry, 1976, 15, 5631–5640.

41. F. Natalio, R. André, A. F. Hartog, B. Stoll, K. P. Jochum, R. Wever and W. Tremel, Nature Nanotechnology, 2012, 7, 530–535.

42. R. André, F. Natálio, M. Humanes, J. Leppin, K. Heinze, R. Wever, H.-C. Schröder, W. E. G. Müller and W. Tremel, Advanced Functional Materials, 2011, 21, 501–509.

43. J. R. Prohaska, Biochimica et Biophysica Acta (BBA) - Enzymology, 1980, 611, 87–98.

44. X. Wang, X. J. Gao, L. Qin, C. Wang, L. Song, Y.-N. Zhou, G. Zhu, W. Cao, S. Lin, L. Zhou, K. Wang, H. Zhang, Z. Jin, P. Wang, X. Gao and H. Wei, Nature Communications, 2019, 10, 704.

45. S. Sarkar, M. Kabir, M. Greenblatt and T. Saha-Dasgupta, Journal of Materials Chemistry A, 2013, 1, 10422–10428.

46. T. Denayer, T. Stöhr and M. Van Roy, New Horizons in Translational Medicine, 2014, 2, 5–11.

47. N. Wang, M. Gray, X.-H. Lu, J. P. Cantle, S. M. Holley, E. Greiner, X. Gu, D. Shirasaki, C. Cepeda, Y. Li, H. Dong, M. S. Levine and X. W. Yang, Nature Medicine, 2014, 20, 536–541.

48. A. Johri and M. F. Beal, Biochimica et Biophysica Acta (BBA) - Molecular Basis of Disease, 2012, 1822, 664–674.

49. P. Kumar, S. S. V. Padi, P. S. Naidu and A. Kumar, Fundamental & Clinical Pharmacology, 2007, 21, 297–306.

50. A. C. Ludolph, F. He, P. S. Spencer, J. Hammerstad and M. Sabri, Canadian Journal of Neurological Sciences 2015, 18, 492–498.

51. R. G. Mealer, S. Subramaniam and S. H. Snyder, The Journal of Neuroscience, 2013, 33, 4206–4210.

52. C. E. Teunissen, H. W. M. Steinbusch, M. Angevaren, M. Appels, C. de Bruijn, J. Prickaerts and J. de Vente, Neuroscience, 2001, 105, 153–167.

53. M. N. Herrera-Mundo, D. Silva-Adaya, P. D. Maldonado, S. Galván-Arzate, L. Andrés-Martínez, V. Pérez-De La Cruz, J. Pedraza-Chaverrí and A. Santamaría, Neuroscience Research, 2006, 56, 39–44.

54. H. H. Yin and B. J. Knowlton, Nature Reviews Neuroscience, 2006, 7, 464–476.

55. Patrick E. Rothwell, Marc V. Fuccillo, S. Maxeiner, Scott J. Hayton, O. Gokce, Byung K. Lim, Stephen C. Fowler, Robert C. Malenka and Thomas C. Südhof, Cell, 2014, 158, 198–212.

56. P. Pla, S. Orvoen, F. Saudou, D. J. David and S. Humbert, Frontiers in Behavioral Neuroscience, 2014, 8.

57. G. K. Shinomol and Muralidhara, NeuroToxicology, 2008, 29, 948–957.

58. R. H. Silva, V. C. Abílio, S. R. Kameda, A. L. Takatsu-Coleman, R. C. Carvalho, R. d. A. Ribeiro, S. Tufik and R. Frussa-Filho, Progress in Neuro-Psychopharmacology and Biological Psychiatry, 2007, 31, 65–70.

59. M. P. Mattson and A. Cheng, Trends in Neurosciences, 2006, 29, 632–639.

