## Supplementary Information for "Incorporation of a Biocompatible Nanozyme in Cellular Antioxidant Enzyme Cascade Reverses Huntington’s Like Disorder in Preclinical Model"

[1]Department of Chemical, Biological and Macromolecular Sciences,
S. N. Bose National Centre for Basic Sciences,
Block JD, Sector 3, Salt Lake, Kolkata 700106, India

[2]Department of Zoology,
Uluberia College, University of Calcutta,
Uluberia, Howrah 711315, India

[3]Department of Zoology,
Vidyasagar University,
Rangamati, Midnapore 721102, India

[4]Department of Microbiology,
St. Xavier's College,
30, Mother Teresa Sarani, Kolkata 700016, India

[5]Technical Research Centre,
S. N. Bose National Centre for Basic Sciences,
Block JD, Sector 3, Salt Lake, Kolkata 700106, India

[6]Research & Development Division,
Dey's Medical Stores (Mfg.) Ltd,
62, Bondel Road, Ballygunge, Kolkata 700019, India

[7]Department of Condensed Matter Physics and Material Sciences,
S. N. Bose National Centre for Basic Sciences,
Block JD, Sector 3, Salt Lake, Kolkata 700106, India

[8]Department of Pathology,
Coochbehar Govt. Medical College and Hospital,
Silver Jubilee Road, Coochbehar 736101, India

[9]Department of Pediatric Medicine,
Nil Ratan Sircar Medical College and Hospital,
138, Acharya Jagadish Chandra Bose Road, Sealdah, Kolkata 700014, India

### Supplementary Tables and Figures

**Table S1: Comparison of kinetic parameters of different GPx mimic.**

| Name of the Enzyme | $K_M$ | $k_{cat}$ | $V_{max}$ | $k_{cat}/K_M$ | pH |
|---|---|---|---|---|---|
| C-$Mn_3O_4$ NP | 0.11 mM | 69.12 $min^{-1}$ | $1.58 \times 10^{-6}$ M $s^{-1}$ | 10530.17 $M^{-1}$ $s^{-1}$ | NA |
| Ebselen | 0.83 mM | 2.36 $min^{-1}$ | $3.13 \times 10^{-6}$ M $s^{-1}$ | 47.5 $M^{-1}$ $s^{-1}$ | NA |
| Ebselen (for GSH) | 2.34 mM | 3.85 $min^{-1}$ | $5.13 \times 10^{-6}$ M $s^{-1}$ | 27.33 $M^{-1}$ $s^{-1}$ | NA |
| $CoFe_2O_4$ NP | 8.89 mM | NA | $1.93 \times 10^{-8}$ M $s^{-1}$ | NA | NA |
| HRP | 3.7 mM | 58 $min^{-1}$ | $8.71 \times 10^{-8}$ M $s^{-1}$ | 15675.68 $M^{-1}$ $s^{-1}$ | 3.5 |
| 6-SeCD | 0.35 mM | 5.5 $min^{-1}$ | NA | 255 $M^{-1}$ $s^{-1}$ | NA |
| 6-diSeCD | 0.243 mM | 10.81 $min^{-1}$ | NA | 740 $M^{-1}$ $s^{-1}$ | NA |
| 2-SeCD | 0.321 mM | 16.0 $min^{-1}$ | NA | 830 $M^{-1}$ $s^{-1}$ | NA |
| GPx (Rabbit liver) | 0.01 mM | 5780 $min^{-1}$ | NA | $9.63 \times 10^{6}$ $M^{-1}$ $s^{-1}$ | NA |
| 2-TeCD | 0.33 mM | 26.65 $min^{-1}$ | $5.33 \times 10^{-7}$ M $s^{-1}$ | 1331.67 $M^{-1}$ $s^{-1}$ | NA |
| $V_2O_5$ Nanowire | 0.11 mM | 3.9 $min^{-1}$ | $7.16 \times 10^{-6}$ M $s^{-1}$ | 590 $M^{-1}$ $s^{-1}$ | 7.4 |
| $Fe_3O_4$ NP | 154 mM | 1430 $min^{-1}$ | $9.78 \times 10^{-8}$ M $s^{-1}$ | 154 $M^{-1}$ $s^{-1}$ | 3.5 |
| $Fe_3O_4$ NP | 54.6 mM | NA | $1.8 \times 10^{-8}$ M $s^{-1}$ | NA | 7.4 |
| Selenoglutaredoxin | 0.01 mM | 0.042 $min^{-1}$ | NA | 70 $M^{-1}$ $s^{-1}$ | NA |
| GPx (Bovine) | 5.3 mM | 9000 $min^{-1}$ | NA | 28301 $M^{-1}$ $s^{-1}$ | NA |
| $MoO_3$ Nanowire | 5.61 mM | NA | $4.6 \times 10^{-6}$ M $s^{-1}$ | NA | NA |

NA: Data not available.

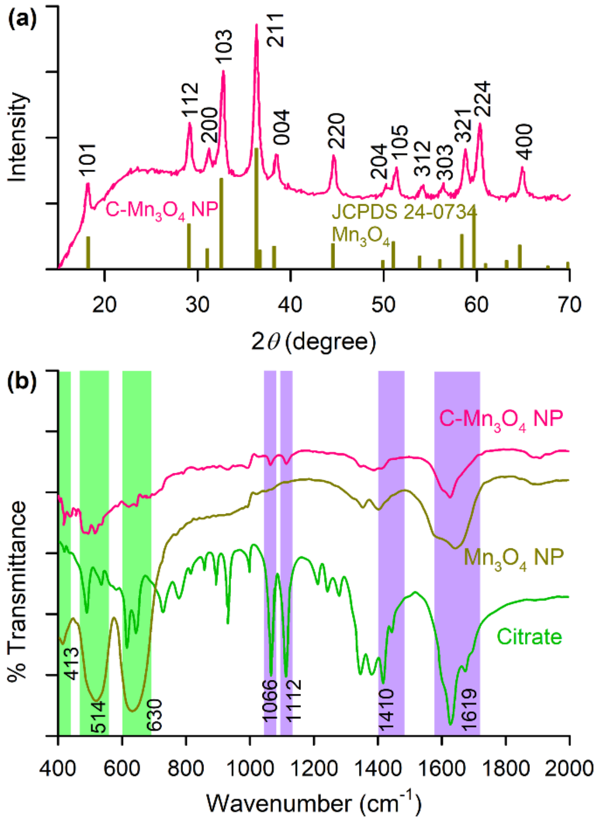


**Supplementary Figure S1. Characterization of C-$Mn_3O_4$ NPs.** (a) XRD and (b) FTIR spectra of C-$Man_3O_4$ NPs.

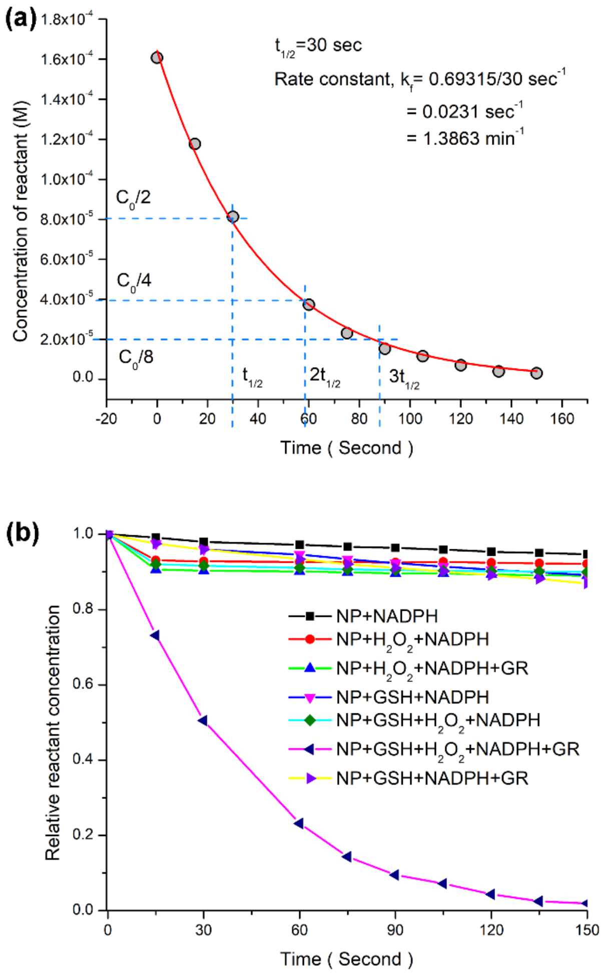


**Supplementary Figure S2.** (a) The enzymatic reaction of $H_2O_2$ catalysis follows first order kinetics. (b) The reaction kinetics at different reaction conditions listed in the legend.

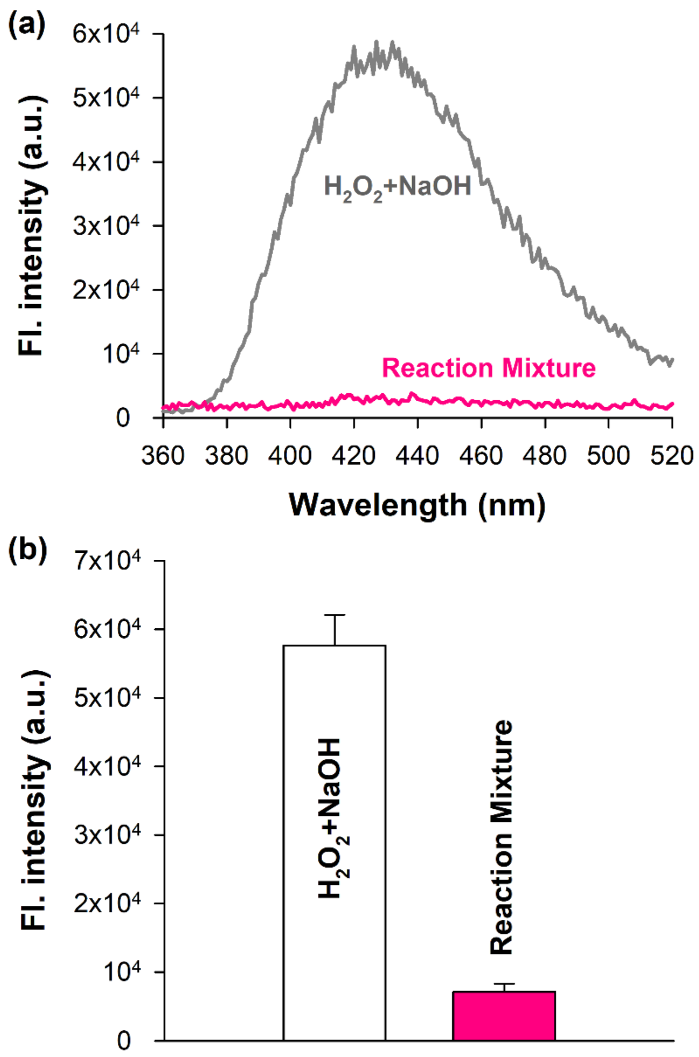


**Supplementary Figure S3. Luminol assay.** (a) The chemiluminescence spectra of luminol in presence of $H_2O_2$ + NaOH (positive control). While in reaction mixture no chemiluminescence peak of luminol was observed. (b) Quantitative estimation of the chemiluminescence intensity at 430 nm.

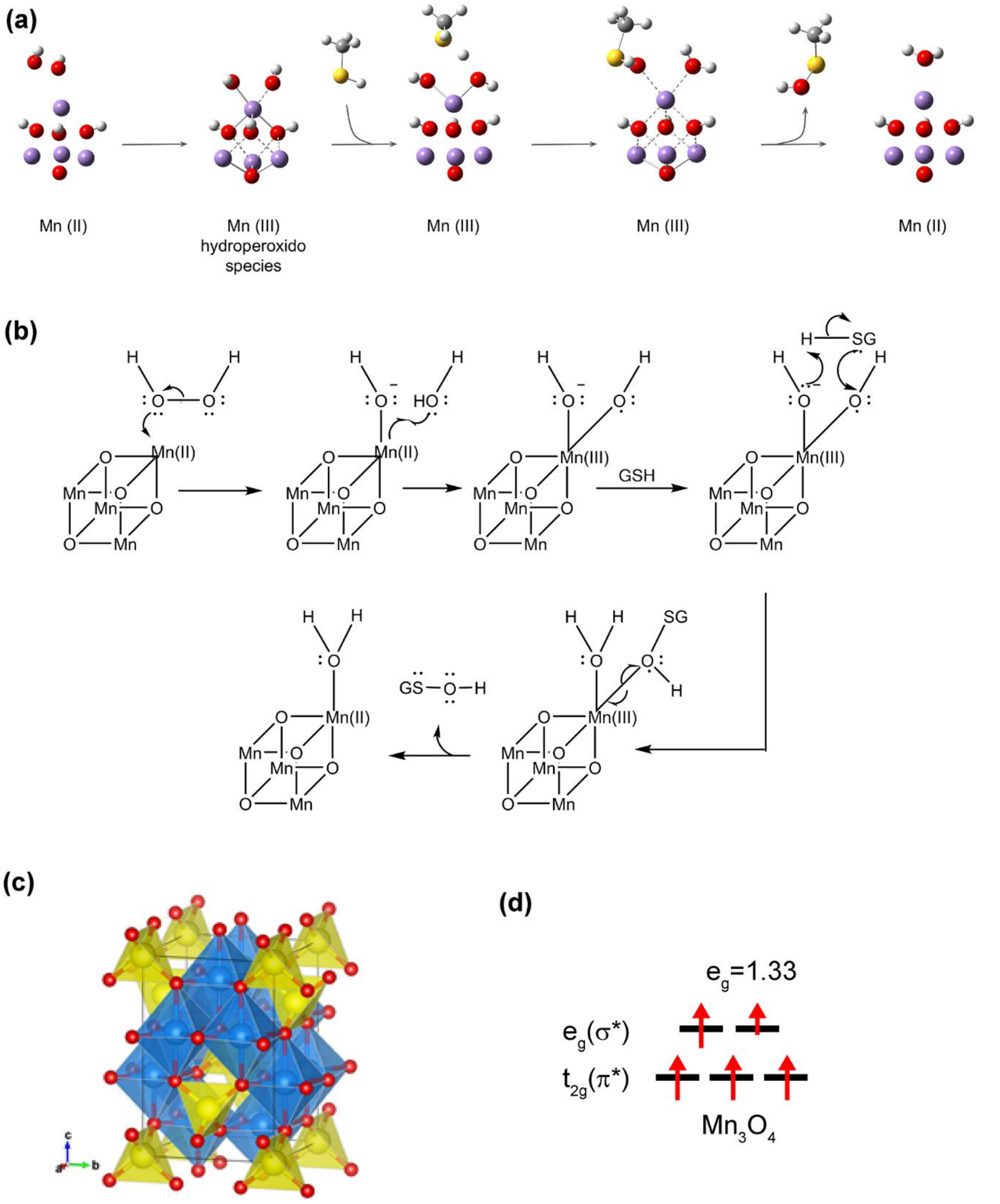


**Supplementary Figure S4. Computational analysis of reaction mechanism.** (a) DFT optimized atomistic model of $H_2O_2$ splitting on the $Mn_4O_4$ cubane complex. Color keys: Mn, purple; O, red; C, gray; S, yellow; H, white. (b) Mechanism of nano-enzymatic action. (b) Peroxidase reaction mechanism. $H_2O_2$ adsorbs on the Mn(II) catalytic center by donating a lone pair from one oxygen atom to a vacant 4sp3 orbital of Mn(II). This leads to the splitting of $H_2O_2$ into two OH groups. In this process one $e_g$ electron of Mn(II) is shared with one of the OH groups, oxidizing the Mn(II) center. Then a proton is transferred from GSH to the OH- attached to the Mn(III) center forming a water and GSattacks the OH group forming GSOH which gets dissociated form the Mn(III) center reducing it to Mn(II). (c) Geometry of $Mn_3O_4$ spinel unit cell with 4 Mn(II), 8 Mn(III) and 16 O atoms. Tetrahedral Mn(II) centers are shown in yellow, Octahedral Mn(III) centers are shown in blue and oxygen atoms are shown in red. (d) Average 3d electron occupancy of $t_{2g}$ ($\pi^*$) and $e_g$ ($\sigma^*$) antibonding orbitals associated with the transition metal Mn.

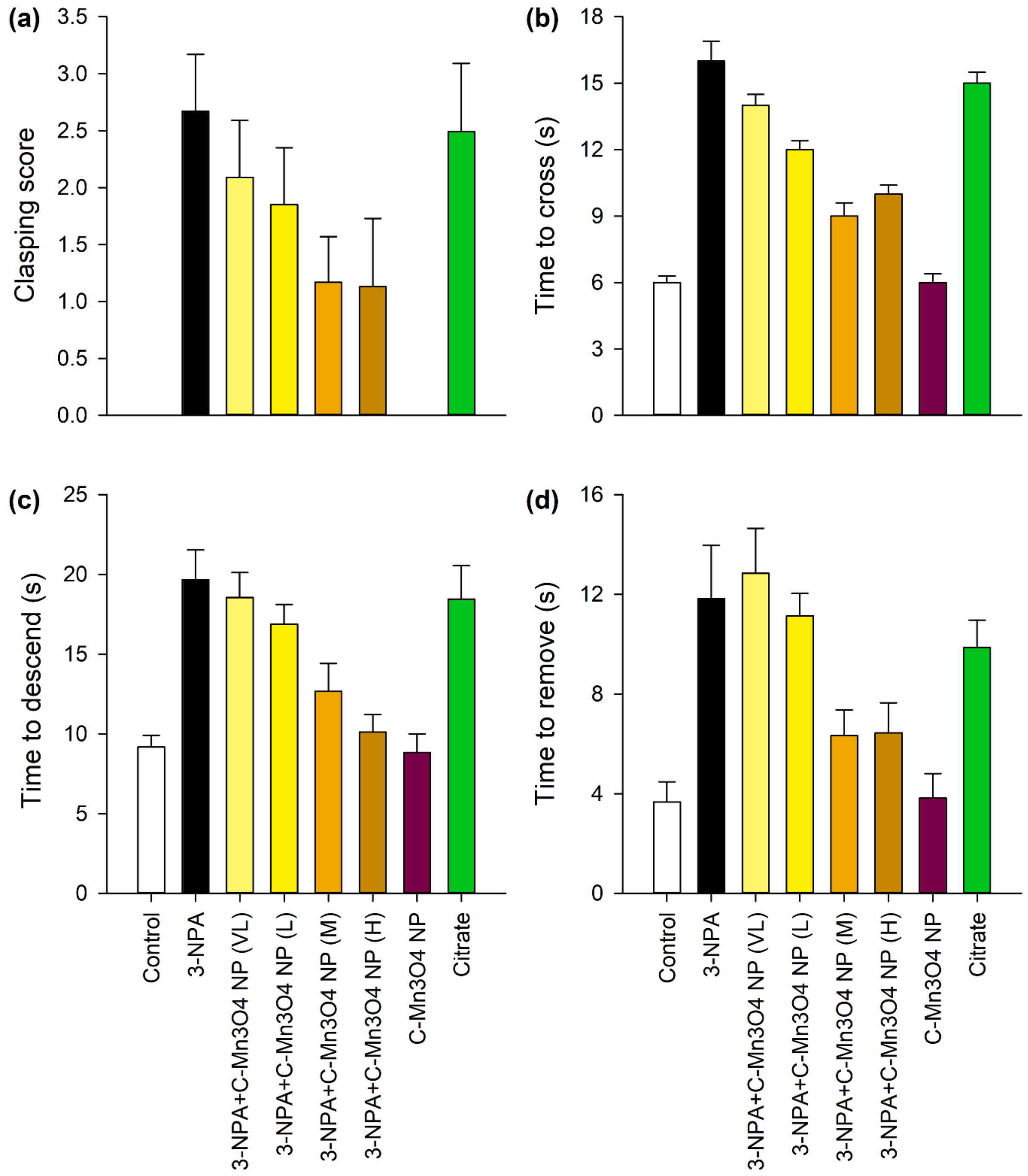


**Supplementary Figure S5. Effect of different doses of C-$Mn_3O_4$ NPs and citrate on 3-NPA induced Huntington's like motor dysfunction.** (a) Hindlimb clasping reflex. (b) Time to cross a beam. (c) Time to descend a pole. (d) Time to remove nasal adhesive.

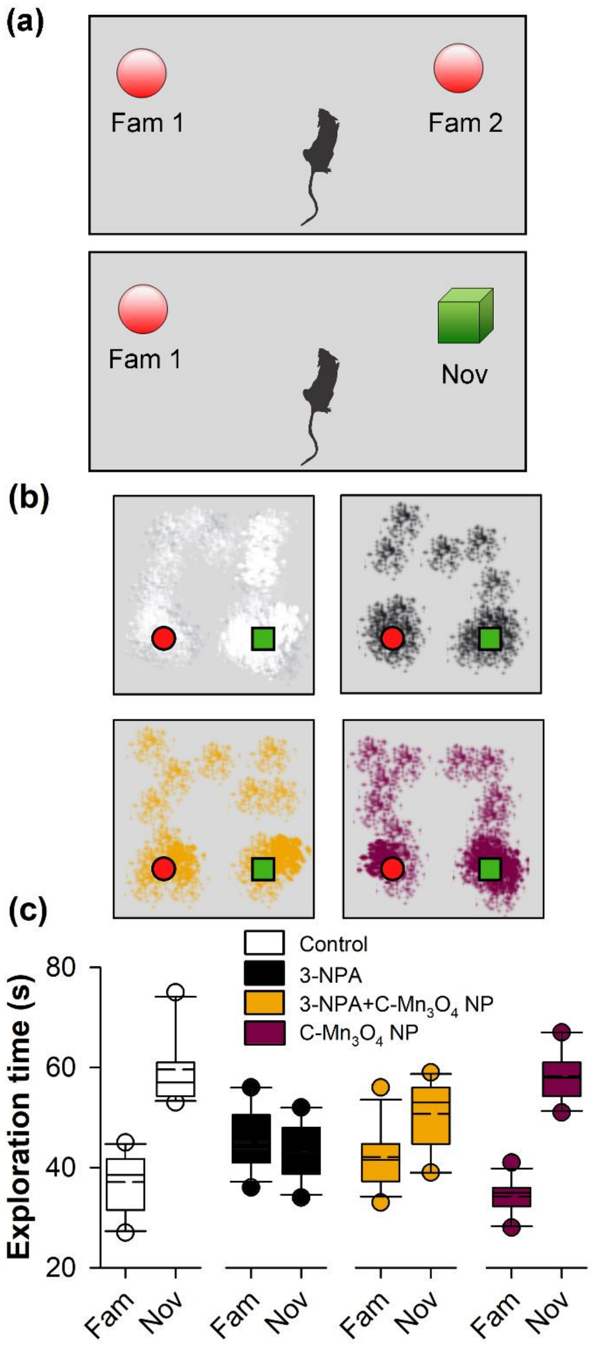


**Supplementary Figure S6. Novel object recognition test.** (a) The experimental setup. (b) The movement pattern of the mice during experiment. (c) Novel object exploration time compared to the familiar one.

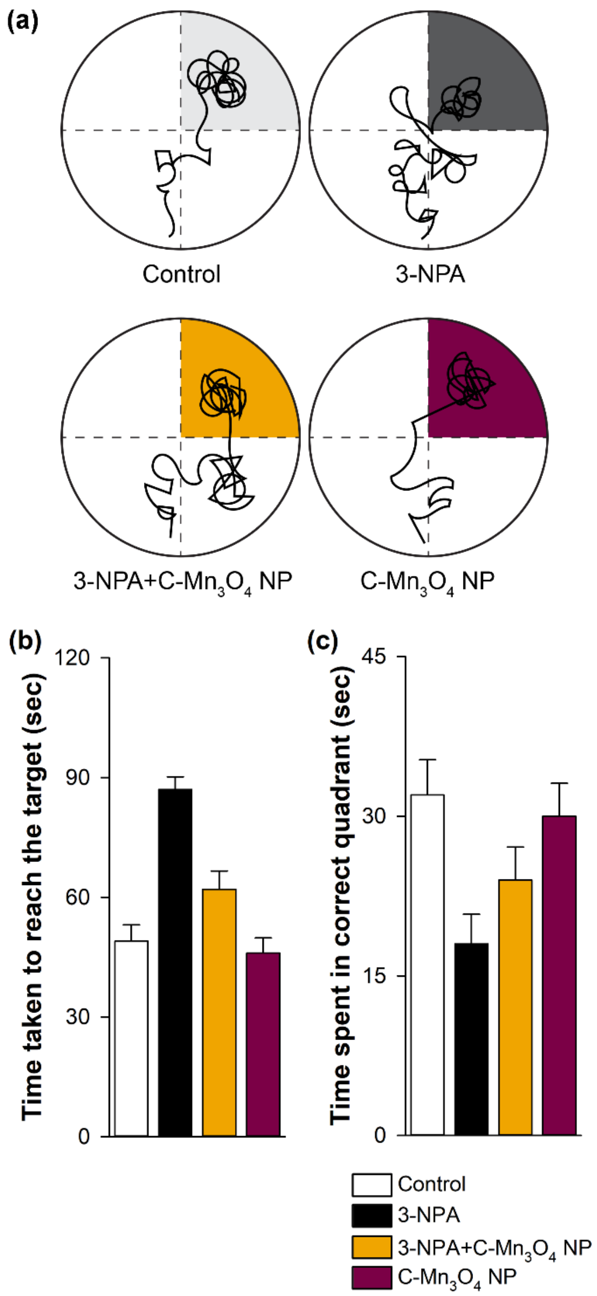


**Supplementary Figure S7. Morris Water Maze Test.** (a) The movement of mice during experiment. The shaded region indicates the target quadrant. (b) Time taken to reach the target. (c) Time spent in the correct quadrant.

### Material and Methods:

##### Synthesis of citrate functionalized $Mn_3O_4$ nanoparticles.

For synthesis of bulk $Mn_3O_4$ nanoparticles at standard temperature and pressure we followed a bottom up approach. In a typical procedure, 0.598 g of $MnCl_2$, $4H_2O$ (3 mmol) was added to 30 ml of ethanol amine (EA) in a beaker, and ultrasonicated at 56 kHz operating frequency for 15 mins. This dissolves $MnCl_2$ to form a clear brown solution. Then equal amount of Mili-Q (from Millipore) water (30 ml) was added and the resultant mixture was stirred at room temperature for 6 hrs. Then the suspension was centrifuged at 3000 rpm for 15 minutes, and the black precipitate was subsequently washed three times using ethanol in order to remove excess EA. After that it was dried in an incubator at 60°C to get a glossy black powder, the as prepared $Mn_3O_4$ NPs.

To functionalize the as-prepared $Mn_3O_4$ NPs with ligand citrate, at first, 0.5 M citrate solution (pH 7.0) was prepared in Milli-Q water. In the ligand solution, as-prepared $Mn_3O_4$ NPs (~150 mg of powder $Mn_3O_4$ NPs in 6 ml ligand solution) were added and extensively mixed for 10 hrs in a cyclomixer. Finally, we filtered out the nonfunctionalized large NPs using a syringe filter (0.22 μm diameter). The resulting filtrated solutions (after proper dilution) were used in further experiments.

##### Characterization of C-$Mn_3O_4$ NPs.

For physicochemical characterization of the NPs various microscopic and spectroscopic studies were performed. For preparation of TEM samples, C-$Mn_3O_4$ NPs were drop casted onto a 300-mesh carbon-coated copper grid and airdried for 12 hrs. TEM micrographs were documented using a FEI TecnaiTF-20 field emission high-resolution transmission electron microscope (FEG-HRTEM) operating at 200 kV. XRD patterns was obtained by employing a scanning rate of 0.02° $s^{-1}$ in the 2θ range from 10° to 80° by PAN analytical XPERT-PRO diffractometer equipped with Cu Kα radiation (at 40 mA and 40 kV). Optical absorbance spectra of the solutions were recorded using a quartz cuvette of 1 cm path length in Shimadzu Model UV-2600 spectrophotometer. For the fluorescence imaging study of liquid NPs, Olympus BX51 fluorescence microscope was used. Jobin Yvon Model Fluoromax-3 Fluorimeter was used to study the characteristic fluorescence excitation and PL spectra of $Mn_3O_4$ NP solutions.

##### GPx-like activity of C-$Mn_3O_4$ NPs.

The GPx-like catalytic activity of C-$Mn_3O_4$ NPs was studied using the GR-coupled assay by following the decrease in the concentration of NADPH spectrophotometrically at 340 nm [1,2] on a Shimadzu Model UV-2600 spectrophotometer under kinetic mode. In a typical assay (1 ml), the reactants were added in the following order, C-$Mn_3O_4$ NPs (1.3 μM), GSH (2mM), NADPH (400 μM), GR (1.7 units), $H_2O_2$ (240 μM) in 100 mM, pH 7.4 phosphate buffer and reaction rate was followed for 30 s at 25°C. The control reactions were performed in the absence of at least one of the reactants. The steady-state kinetics of C-$Mn_3O_4$ NPs were studied by varying the concentration of C-$Mn_3O_4$ NPs, $H_2O_2$, and GSH at a time and at the fixed concentration of GR and NADPH in 100 mM phosphate buffer (pH 7.4). Michaelis–Menten curves and Lineweaver–Burk plots were obtained by using origin 9.0 software. To understand the effect of haloperoxidase substrates on the

GPx-like activity of C-$Mn_3O_4$ NPs, the GPx-like activity of C-$Mn_3O_4$ NPs was followed in the presence of KI (2 mM) and tyrosine or dopamine (2 mM) with GR-coupled assay in 100 mM phosphate buffer (pH 7.4).

**Computational study.**

Computational study was performed on the $Mn_4O_4$ cluster using density functional theory (DFT) as implemented in Gaussian 16 software. Geometry optimizations were performed with unrestricted B3LYP functional and Pople's double zeta basis set 6-31g with added diffuse and polarization functions on the heavy atoms. High spin geometry of Mn(II) was considered. The spin multiplicity of the system was 21. Three oxygen atoms were passivated with three hydrogens, giving rise to +3 charge. GSH was mimicked with RSH, where the R is a methyl group. The first step of the reaction (Figure 2g) was modeled by O--O bond length perturbation of adsorbed $H_2O_2$. The proton transfer step was modeled by distance perturbation of donor hydrogen and acceptor oxygen. RSOH formation was modeled by sulfur-oxygen distance perturbation.

The $e_g$ occupancy presented here is the average between integer occupations of Mn(II) and Mn(III) metal centers as suggested by Wang *et al.* There are no fractional electrons occupying these orbitals.

**Animals and Treatment.**

The selected animals (C57BL/6j mice) of age 6-8 weeks were transferred to a single housed room for acclimatization 4 weeks prior to tests.

The treatments were as follows,

Group 1: Control (Normal saline 16 days).

Group 2: 3-Nitropropionic acid treated (10 mg $kg^{-1}$ body weight (BW) for alternative 4 days).

Group 3: 3-Nitropropionic acid (10 mg $kg^{-1}$ body weight (BW) for alternatively first 4 days days) + C-$Mn_3O_4$ NPs (5 mg $kg^{-1}$ (BW) for 16 days).

Group 4: 3-Nitropropionic acid (10 mg $kg^{-1}$ body weight (BW) for alternatively first 4 days) + Citrate (5 mg $kg^{-1}$ (BW) for 16 days).

Group 5: C-$Mn_3O_4$ NPs (5 mg $kg^{-1}$ (BW) for 16 days).

Necessary ethical permissions were taken from Institutional Animal Ethics Committee. All tests were performed following the guide line of CPCSEA, India.

**Behavioral assays**

All mice were subjected to sequential tests started with open field test then followed by tail flick, rotarod, beam walking, elevated plus maze, light and dark box, forced swim test, Sucrose preference test, pole test, novel object recognition and Morris water maze. Some tests like open field test, hind-limb clasping test, beam walking, elevated plus maze were performed in an interval of 2 days. Other Tests were done before treatment and at the end of 16$^{th}$ day. Between two consecutive tests there were a period of time of hour in which the animals left undisturbed.

**Hindlimb Clasping Test:** This study is normally used to check the disease progression in Huntington's disease. A well-studied method is followed to carry out this experiment in every alternative days up to $16^{th}$ days [3].

**Open field test (OFT):** This test measures thigmotaxis behavior (to measure the anxity) and motor deflects of the subject. For this test a box, divided into a center and peripheral zone, was used (46cm X 46cm X 30cm). Each animal was placed into the box and allowed to explore the box for 5 minutes and an automated video camera was used to record its behavior. Time spent by the animal in the central zone and peripheral zone was measured to evaluate its anxiety level and motor movement.

**Tail flick test:** Tail-flick test is a useful study based on the measurement of the latency of the avoidance response to thermal stimulus in rodents. Basically, a thermal stimulus is applied to the tail; when the animal feels discomfort, it reacts by a sudden tail movement. The tail flick reaction time is recorded and indicated as an index of animal pain sensitivity[4].

**Rotarod:** Motor skill learning and motor coordination measured to assess the fine motor deflects in rodents [5]. We have trained the mice for three days before the main tests. In training they were placed on the rod and rotation speed was set at 10 rpm. In the training day the mice were placed to the rotarod for 2 mins and the latency to fall was recorded. Each mice was subjected for 5 consecutive trails.

**Beam walking:** This test is a useful way to access fine motor coordination and balance of rodents. Each mouse was placed at the end of a narrow beam of diameter 2.5 cm and 50 cm long which it had to cross. At the starting end an aversive stimuli, i.e. a brightly lit blub, was given and at the opposite end a platform was placed into darkness where some food pellets were kept as rewards. Each animal was evaluated by measuring the delay period before starting, the time to reach into the platform or fall or false start, the distance covered by the animal before falling from the beam and the number of paw slip. Each mice were given 5 trails.

**Elevated plus maze:** Elevated plus maze is widely used for evaluating anxiety like behavior in the rodents. The test apparatus is made up of 4 arms two of which is closed and the other two is open. The apparatus is placed 40 cm above the ground level and the closed and open arms were perpendicularly crossed each other in a central area. All the arms were brightly lit. Closed arms were covered by wall of 15 cm and open arms had a barrier of 2 cm that prevent slipping of animal.

Animals were scored depending on the entry to the open arm and time spent in it. This score reflect the measurement of anxiety that is induced in the animal by the open space.

**Light and Dark Box:** The light/dark test is established on the natural aversion of rodents to brightly illuminated areas and also on their spontaneous exploratory behavior in response to mild stressors like novel environment or light. Here the test apparatus is divide in two compartment, light area (aversion area) is two third of the apparatus and the dark area (safe area) is one third. Each mice were tested for 5 minutes. The time spent in both areas and the number of transit between two areas were measured.

**Forced Swim test:** Individual mouse was placed into a glass beaker of diameter 16 cm and height of 26 cm. The apparatus was filled up to 15cm with water of temperature $25^0$C. Time spent by the animal in the water was recorded and immobility (refers to the minimal movement of the animal to keep its head over the water surface) of the animal was measured. An increased in immobility reflects increase in depression. After test all animals were dried properly and put into new cages.

**Sucrose preference test:** Consumption of a 2% sucrose solution between 08:00 am and 09:00 pm was recorded in all mice to measure sucrose anhedonia[6]. The tests were done by following a previously standardise protocol [7].

**Morris water maze:** After acclimatization, animals from each of the groups were taken for the Morris water maze study to evaluate their spatial learning ability. The complete study consists of training period of 7 days before starting the treatment and then test period. This training was done in a pool with a platform (target) whose top was lifted above the water surface. Distal cues were clearly visible for the animals at the time of training. The animals were placed in maze in a random fashions from 4 different start points. On 16th days of treatment animals were evaluate for their spatial learning ability. All the animals were placed from a fixed point and the platform were hidden beneath the water surface in this session. All the animals were given 90 sec for exploration. Time taken to reach the platform and no of time the animal entered in the target quadrant was analysis for accessing their spatial learning and memory that give insight to hippocampal damage.

**Novel object recognition test:** For the evaluation of memory alteration among the four experimental groups we have employed novel object recognition test following a standard protocol used in pervious study [7].

**Serum isolation.**

At the end of the experimental period, the animals were euthanized and decapitated after being fasted. Blood was collected from retro orbital plexus just before sacrifice, kept in sterile non-heparinized tubes in slanting position for 45 min and centrifuged at 3500×g for 20 min. The clear serum was obtained and used in subsequent biochemical analysis.

**Histopathological examination.**

For microscopic evaluation, a conventional technique of paraffin wax sectioning and differential staining was used [8]. 2 h, dehydrated in graduated ethanol (50–100%), cleared in xylene, and embedded in paraffin. Microtome was used to prepare ultrathin sections (4–5 μm), followed by staining with hematoxylin and eosin (H&E) and silver stain. Histopathological changes were examined under the microscope (Olympus BX51) equipped with a CCD based camera.

**Tissue homogenate preparation.**

Tissue samples were collected, homogenized in cold 0.1 mM phosphate buffered saline (PBS; pH 7.4), and centrifuged at 10,000 rpm at 4°C for 15 min. The supernatants were collected to determine the activity of SOD, CAT, GPx and GSH as well as the content of malondialdehyde (MDA).

**Assessment of lipid peroxidation & antioxidant enzyme activity.**

The supernatants were used to determine the activity of SOD, CAT, GPx and GSH as well as the content of MDA. SOD, CAT and GSH activities were estimated using commercially available test kits (Sigma-Aldrich, MO, USA) following the protocols described by the manufacturer. To assess the extent of lipid peroxidation, the level of malonyldialdehyde (MDA), a substance that reacts with thiobarbituric acid, was determined in the homogenates of organs and in serum according to the method of Buege[9].

**Mitochondria Isolation.**

Mitochondria isolation was done from mouse brain following the method of Graham[10] with some modifications. In brief, brains were excised and homogenized in brain homogenization medium that contains 250 mM D-mannitol, 125 mM sucrose, 0.05 mM EGTA, 0.01% BSA, 10 mM HEPES (pH 7.2), 1x protease inhibitors. Then the homogenates were centrifuged for 15 minutes at 700 x g and the supernatants were again centrifuged as the previous step. Then supernatant were washed collected and centrifuged at 10000 x g for 15 minutes. The resultant pellets were dissolved in ice cold buffer with added digitonin. Again it was centrifuges at 10000 x g for 15 minutes and discared the supernatant while re-suspended the pellet in extraction buffer. For all procedure temperature was maintained at 4°C. Commercially available kit (Autospan Liquid Gold, Span Diagnostics Ltd., India) was used for determining protein concentration following the protocol described by the manufacturer.

**Complex IV (of respiratory chain) activity.**

Total complex IV activity was measured spectrophotometrically using isolated mitochondria [11]. Briefly, reduced cytochrome c was prepared by mixing cytochrome c and ascorbic acid in potassium phosphate buffer. Complex IV activity was taken as the rate of ferrocytochrome c oxidation to ferricytochrome c, detected as the decrease in absorbance at 550 nm.

**Measurement of mitochondrial membrane permeability transition**.

Opening of the pore causes mitochondrial swelling, which results in reduction of absorbance at 540 nm. Mitochondrial permeability transition (swelling assay) was monitored as changes at 540 nm at 10 s intervals over 10 min time with 250 µg mitochondrial protein in the swelling buffer, which contained 120 mM KCl (pH 7.4) and 5 mM $KH_2PO_4$ .

**Measurement of mitochondrial membrane potential.**

The mitochondrial membrane potential (ΔΨm) was measured using the fluorescent probe rhodamine 123 (Sigma) [12, 13]. Because rhodamine 123 is a cationic dye, it accumulates in the mitochondria driven by ΔΨm. Under appropriate loading conditions, the concentration of rhodamine 123 within the mitochondria reaches sufficiently high levels that it quenches its own fluorescence (λex = 503 nm, λem = 527 nm). If the mitochondria depolarize, rhodamine 123 leaks out into the cytoplasm and is associated with a reduction in the amount of quenching. Thus the changes in ΔΨm are revealed as changes in total fluorescence intensity following the method of Chen[14].

**Mitochondrial dehydrogenases activity.**

The methyl tetrazolium (MTT) assay was used as a colorimetric method for the estimation of mitochondrial dehydrogenases activity[15, 16] in isolated brain mitochondria. Briefly, mitochondrial suspension (1 mg protein/mL) in a buffer containing 320 mM sucrose, 10 mM Tris-HCl, 1 mM EDTA, and pH= 7.4, was incubated with 40 µL of theMTT solution (0.4% w: v) at 37°C (30 min, in the dark). The product of purple formazan crystals was dissolved in DMSO (1 ml). Then, 100 µl of dissolved formazan product was added to a 96 well plate, and the optical density (OD) was measured at λ= 570 nm using an EPOCH plate reader (BioTek Instruments, USA).

**Mitochondrial ATP level.**

A luciferase-luciferin-based kit from Promega (ENLITEN, Madison, WI, USA) was used to assess brain mitochondrial ATP content. Samples and buffer solutions were prepared based on the kit instructions. Briefly, mitochondria fractions (1 mg protein/ml) were treated with 100 µL of trichloroacetic acid (0.5% w: v). Samples were centrifuged (15,000 g, 15 min, 4°C). Then, 100 µL of the supernatant was added to 100 µL of the kit content, and the luminescence intensity was measured at λ= 560 nm.

**Statistical analysis.**

All quantitative data are expressed as mean ± SD unless otherwise stated. One-way analysis of variance (ANOVA) followed by Tukey's multiple comparison test was executed for comparison of different parameters between the groups using a computer program GraphPad Prism (version 5.00 for Windows), GraphPad Software (CA, USA). $p < 0.05$ was considered significant.

#### Reference:


1. N. Singh, M. A. Savanur, S. Srivastava, P. D'Silva and G. Mugesh, *Angewandte Chemie*, 2017, **129**, 14455-14459.

2. S. Ghosh, P. Roy, N. Karmodak, E. D. Jemmis and G. Mugesh, *Angewandte Chemie*, 2018, **130**, 4600-4605.

3. S. J. Guyenet, S. A. Furrer, V. M. Damian, T. D. Baughan, A. R. La Spada and G. A. Garden, *JoVE (Journal of Visualized Experiments)*, 2010, e1787.

4. S.-L. Chen, H.-I. Ma, J.-M. Han, R.-B. Lu, P.-L. Tao, P.-Y. Law and H. H. Loh, *Journal of Biomedical Science*, 2010, **17**, 28.

5. Y. Qian, M. Chen, H. Forssberg and R. Diaz Heijtz, *Genes, Brain and Behavior*, 2013, **12**, 604-614.

6. L. K. Fonken, E. Kitsmiller, L. Smale and R. J. Nelson, *Journal of Biological Rhythms*, 2012, **27**, 319-327.

7. A. Adhikari, M. Das, S. Mondal, S. Darbar, A. K. Das, S. S. Bhattacharya, D. Pal and S. K. Pal, *Biomaterials science*, 2019, **7**, 4491-4502.

8. A. Adhikari, S. Darbar, T. Chatterjee, M. Das, N. Polley, M. Bhattacharyya, S. Bhattacharya, D. Pal and S. K. Pal, *ACS omega*, 2018, **3**, 15975-15987.

9. J. A. Buege and S. D. Aust, in *Methods in enzymology*, Elsevier, 1978, vol. 52, pp. 302-310.

10. J. M. Graham, in *Biomembrane protocols*, Springer, 1993, pp. 29-40.

11. M. Spinazzi, A. Casarin, V. Pertegato, L. Salviati and C. Angelini, *Nature Protocols*, 2012, **7**, 1235.

12. R. C. Scaduto Jr and L. W. Grotyohann, *Biophysical journal*, 1999, **76**, 469-477.

13. J. D. Ly, D. R. Grubb and A. Lawen, *Apoptosis*, 2003, **8**, 115-128.

14. L. B. Chen, *Annual review of cell biology*, 1988, **4**, 155-181.

15. T. Mosmann, *Journal of immunological methods*, 1983, **65**, 55-63.

16. M. M. Ommati, R. Heidari, V. Ghanbarinejad, A. Aminian, N. Abdoli and H. Niknahad, *Nutritional Neuroscience*, 2018, 1-13.
